# MisTIC: Missegmented Transcript Inference Correction for Improved Spatial Transcriptomics Analysis

**DOI:** 10.64898/2025.12.11.693759

**Authors:** Yuqiu Yang, Erica DePasquale, David Adeleke, Xiangfei Xie, Zeyu Lu, Michael D. Nelson, Guanghua Xiao, Xinlei Wang, Yunguan Wang

## Abstract

Imaging-based spatially resolved transcriptomics (SRT) technologies, such as 10X Xenium, MERSCOPE, and CosMx, have revolutionized our ability to study gene expression within the spatial context of tissues at single-cell resolution. The acquisition of such data relies heavily on cell segmentation algorithms, which often produce imperfect boundaries, leading to transcript misassignment. These misassignments can significantly affect downstream analyses, including cell type identification, differential expression analysis, cell-cell communication, and RNA localization. We present MisTIC (Missegmented Transcript Inference Correction), a variational Bayesian model designed to correct transcript misassignment errors without requiring resegmentation. In benchmarking analyses using synthetic data with simulated transcript misassignment, MisTIC demonstrated high sensitivity and specificity in removing misassigned transcripts. In real data applications, MisTIC effectively enhances cell type identification, reduces ambiguity in differential expression analysis, and improves the detection of cell-cell communication. Furthermore, RNA localization analysis based on data corrected by MisTIC revealed that, in T cells located near cancer-associated fibroblasts compared to those farther away, genes involved in T cell activation, inflammation, and cytotoxicity were depleted from cytoplasmic regions despite not being differentially expressed between the two T cell subsets. In conclusion, MisTIC is a powerful tool for correcting transcript misassignment in SRT data. It not only improves the accuracy of routine analyses but also enables novel investigations that provide deeper insights into the dynamics of gene expression.

## Introduction

Spatially resolved transcriptomics (SRT) has emerged as a suite of transformative technologies in molecular biology, allowing biologists to study gene expression while preserving the spatial context of tissues (1). Recent advancements in this omics technology have enabled quantification of RNA molecules at subcellular resolution, which allows researchers to study cellular heterogeneity (2), identify spatial gene expression patterns (3), and characterize tissue organization (4) in unprecedented detail.

In general, SRT technologies can be categorized into two groups depending on the underlying techniques they leverage (1). Sequencing-based SRTs such as Visium HD (5), Seq-Scope (6), and Stereo-seq (7) utilize next-generation sequencing and DNA barcoding to measure the gene expression levels for nearly the whole-transcriptome with a resolution of 0.2-2µ𝑚. These technologies rely on the usage of “spots” which are barcoded grids covering the tissue to provide aggregated expression measurement at the spot level. On the other hand, imaging-based SRTs such as the MERFISH (8) and MERSCOPE from VizGen (9), CosMx and CosMx WTX from NanoString (10, 11), and Xenium from 10X Genomics(12) are based on fluorescence *in situ* hybridization (FISH) to detect hundreds to thousands of genes. Although challenges such as low gene-coverage and sparsity in data (13) remains for these platforms, they possess a key advantage over the other category by providing the localization of individual RNA molecules and cells, which empowers single-cell level analysis and is valuable in generating new insights into intercellular crosstalk, gene regulation and transcriptional dynamics (1).

The analytical pipeline for imaging-based SRT data relies critically on accurate cell segmentation to assign detected transcripts to their cells of origin. This process is often accomplished by incorporating nuclear staining such as DAPI (14) which binds to AT-rich regions of DNA, providing clear visualization of cell nuclei that serve as anchors for defining each individual cell. Cellular boundaries are then further delineated by membrane staining or computational methods such as Watershed (15, 16), Cellpose (17), Baysor (18), FICTURE (19), BIDCell (20), amongst others.

Despite the numerous cell segmentation algorithms proposed, accurate cell segmentation still presents significant technical challenges (21–23), particularly in densely packed tissues where cell boundaries are difficult to discern. Furthermore, irregular morphologies of cells, imaging artifacts, and staining inconsistencies can further complicate accurate cellular boundary detection. These factors inevitably lead to segmentation errors that propagate through downstream analyses. One of the most direct consequences of inaccurate segmentation is the misassignment of transcripts to incorrect cells. These misassigned transcripts may originate from neighboring cells with either similar or vastly different gene expression profiles—the latter posing a more serious problem. This leads to systematically biased gene expression measurements, confounded cell type identification, and obscured genuine spatial patterns. Such errors frequently manifest as the co-expression of mutually exclusive genes within the same cell and ambiguity in differential expression analyses. While post-hoc corrections using domain-specific knowledge can partially mitigate these effects, they do not address the root cause. More critically, current cell segmentation algorithms for image-based SRT data lack mechanisms to quantify the confidence of predicted segmentation masks or estimate the fraction of misassigned transcripts within each cell.

Current approaches for addressing segmentation-related errors in image-based SRT data remain limited, with post-processing methods for rectifying errors receiving little attention. To our best knowledge, only two methods are specifically designed for this purpose: ResolVI (24) and FastReseg (25). ResolVI is a VAE-based model that decomposes cell-level count into three components: true expression, neighbor-derived expression, and background noise. However, ResolVI does not reassign individual transcripts. Instead, it only modifies the aggregated cell-by-gene count matrix, limiting its utility for applications sensitive to transcript localization. In contrast, although FastReseg operates at the molecule level, it relies on a sequence of heuristic decision rules to reassign transcripts, limiting its robustness and generalizability across tissues with complex morphology or diverse transcript densities. To address these gaps, we developed MisTIC (**Mis**segmented **T**ranscript **I**nference **C**orrection), a probabilistic model designed to correct transcript misassignment errors without requiring resegmentation. MisTIC leverages the spatial distribution of transcripts and patterns of transcriptional profiles from different cell groups to identify and correct misassigned transcripts. By utilizing the recent advancements in variational Bayes (26, 27), MisTIC is capable of scaling up to tens of millions to hundreds of millions of detected transcripts and provides a probability of misassignment for each individual RNA. Through extensive analyses using both simulated and real imaging-based SRT data, we demonstrate that MisTIC recovers the true transcriptional profile of cells with high accuracy, enhances the detection of cell type-specific markers, improves the resolution of cell-cell communication patterns, and facilitates studies on RNA localization.

Overall, MisTIC is effective in detecting erroneously assigned transcripts, is computationally efficient, and can be seamlessly integrated into existing spatial transcriptomics analysis pipelines. We believe that MisTIC represents a significant advancement in addressing a critical limitation of imaging-based SRT technologies. MisTIC was tested in a wide range of imaging-based SRT datasets including Xenium/Xenium Prime, MERSCOPE, and CosMx WTX, and it decreased ambiguity in both cell type assignment and differentially expressed genes, reduced false positives and false negatives in CCC predictions, and uncovered biological insights otherwise masked by misassigned transcripts. Currently, MisTIC is available as an open-source project at https://github.com/yunguan-wang/MisTIC-Wanglab.

## Results

### Erroneous cellular boundaries can obscure biological signals

We first demonstrated the existence of extensive transcript misassignment and its impact on gene expression profiles of the cells. We evaluated imaging-based SRT datasets profiling lung adenocarcinoma (LUAD) tissues from two commonly used commercial platforms (Fig. S1a and b), 10X Xenium and VizGen MERSCOPE (See Data Availability for details), in the context of bulk RNA-seq data from The Cancer Genome Atlas Program (TCGA) LUAD dataset (n=452), and reference scRNA-seq data (28) from LUAD patients. Through analysis of the reference scRNA-seq data using only genes included in the Xenium SRT data, we identified marker genes for major cell types in LUAD, including tumor cells, T cells, B cells, fibroblasts, and macrophages. Expression of these genes were largely exclusive (Fig. 1a), however, in the SRT data some of these genes were expressed in non-specific cell types at low, but not negligible levels (Fig. 1b). For example, we found 35.4% B cells expressed *CD3E,* and 29.5% Macrophages expressed *EPCAM*. Upon examining the localization of genes expressed in non-specific cell types, we found they were enriched in cells in close contact with another cell type, and close to the predicted cell borders (Fig. 1c). These results demonstrated that the misassignment of transcripts adds heterotypic genes to a cell’s transcriptome profile and leads to artificial cell states similar to the “doublet” cells from scRNA-seq data.

**Fig. 1.**
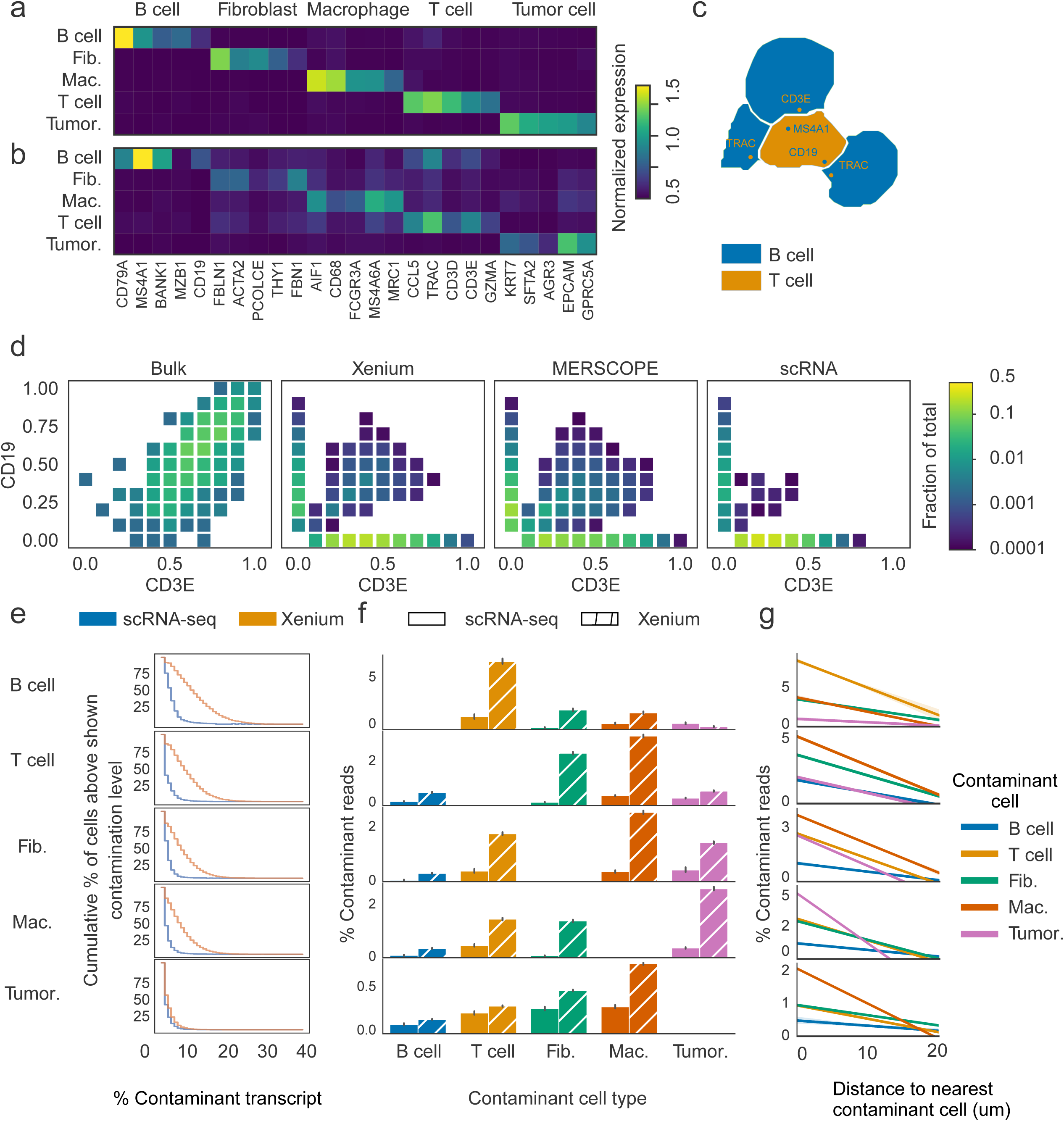
Erroneous cellular boundaries can obscure biological signals. (a, b) Average expression of marker genes in cell types from the reference scRNA-seq (a) and the Xenium SRT data (b) from LUAD tissues. Gene expression was normalized by library size and log-transformed. (c) Representative cell masks of B and T cells showing misassigned CD19, MS4A1, CD3E, and TRAC transcripts. (d) Scatter heatmap of CD19 and CD3E in cells expressing at least one of these genes. Each box represents an expression bin defined by CD19 and CD3E expression levels, with bin colors mapped logarithmically to the fraction of cells. (e) Cumulative percentage of contaminant transcripts relative to total transcript counts in different cell types, calculated from reference scRNA-seq (blue) and SRT (orange) data. X-axis shows the percentage of contaminating transcripts, and Y-axis shows the cumulative percentage of cells with at least the contamination level indicated on the X-axis. (f) Contribution of contaminant transcripts by source. Percentages were calculated per cell in scRNA-seq (blank bars) and SRT (slashed bars). Error bars represent standard deviation. (g) Percentage of contaminant transcripts in each cell originating from other cell types as a function of the distance to contaminating cells, where cell distance was defined as the centroid-to-centroid distance.

Next, we evaluated the co-expression of *CD3E* and *CD19*, which are typically mutually exclusive markers of T cells and B cells, respectively, within the same cells in SRT data. We then compared the resulting co-expression patterns with those derived from bulk RNA-seq and scRNA-seq data, which served as controls. In this analysis, we discretized cells into 100 bins defined by pairwise combinations of 10 expression intervals for each of two genes. For each bin, we computed the fraction of total cells it contained. Most of the samples from bulk RNA-seq data co-expressed *CD19* and *CD3E*, which is expected as B and T cells are commonly found in tertiary lymphoid structures in various cancers (29–34). In contrast, co-expression between these two pairs of genes in single cells from the reference LUAD scRNA-seq dataset (35) was very rare. The level of co-expression in cells from SRT data was between those from sc- and bulk RNA-seq data. While the majority of cells fell into bins corresponding to the expression of only *CD19* or *CD3E*, we observed a broad distribution of bins—each containing 0.1-1% of cells—representing moderate co-expression of both genes (Fig. 1d). Despite their individually small fractions, these co-expressing bins were numerous in both assessed SRT datasets. We extended this analysis to additional pairs of mutually exclusive marker genes representing fibroblast/T cells, macrophage/T cells, and macrophage/tumor cells, and observed co-expression within the same cells at the similar level (Fig. S2).

Next, we assessed transcript contamination at the single-cell level in SRT data, using scRNA-seq data as a control. To ensure specificity, we used a conservative marker gene set comprising only the top five selectively expressed genes for each cell type. We then calculated the fraction of transcripts derived from contaminating cell types as a proportion of total transcript counts. Cells from SRT data exhibited higher levels of contamination compared to their scRNA-seq-derived counterparts. For example, among B cells, T cells, fibroblasts, and macrophages in the scSRT data, 47.0% to 73.4% of cells had more than 5% of their transcriptome composed of contaminating genes, whereas only 3.0% to 10.3% of cells in scRNA-seq data showed such contamination (Fig. 1e). A high-degree of contamination, however, was rare: only 0.2% to 2.9% of these cells had at least 25% of their transcripts originating from other cell types.

To clarify the source of these contaminating transcripts, we examined their cell type of origin. In SRT data, the contaminating transcript sources varied substantially across cell types, a pattern that was much less pronounced in scRNA-seq data. For instance, in B cells, most contaminating reads originated from T cells, whereas in T cells, the majority of contamination came from fibroblasts and macrophages (Fig. 1f). These patterns are consistent with known tissue architecture: B and T cells often reside in close proximity within tertiary lymphoid structures in tumors (36–39), and cancer-associated fibroblasts and macrophages are known to regulate T cell activity (40–45). This observation suggested the level of transcript contamination in a cell was influenced by its neighboring cells. To evaluate this possibility, we calculated each cell’s contamination fraction and its distance to the nearest heterotypic cells (of each type) within a 20 µm radius. We found contamination from heterotypic transcripts was negatively correlated with distance and declined to near-zero levels when cells were sufficiently separated (Fig. 1g). In summary, these results indicate that transcript contamination in imaging-based SRT data is frequent and generally contributes only a modest fraction of each cell’s transcriptome, with its extent strongly dependent on spatial proximity to neighboring heterotypic cells.

### MisTIC model overview

To correct for cell segmentation errors that result in misassigned transcripts in imaging-based SRT, we developed a probabilistic framework, MisTIC, for transcript reassignment based on biologically and spatially motivated features. MisTIC operates at the level of individual transcripts and considers the likelihood of reassignment based on three key factors: spatial proximity, gene expression compatibility, and neighborhood expression support (Fig. 2a).

**Fig. 2.**
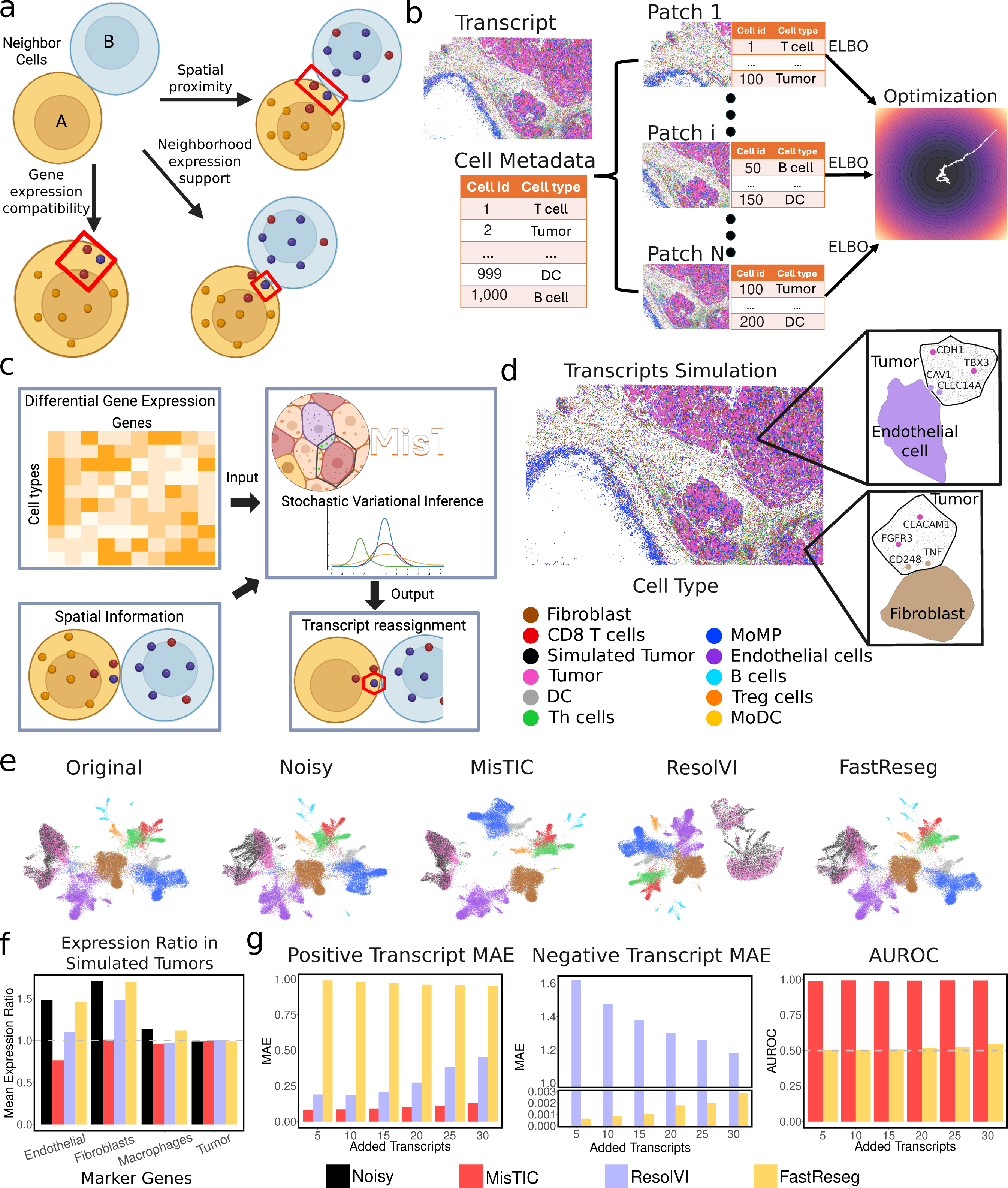
MisTIC model overview and synthetic data results. (a) Three key factors that MisTIC uses for computing the likelihood of reassignment. (b) The training process of MisTIC. (c) Overview of the workflow of MisTIC. (d) Illustration of simulated tumor cells with added transcripts. (e) UMAP embeddings of original, uncorrected, and corrected SRT data (FastReseg, ResolVI, MisTIC). (f) Expression ratios of marker gene expression with the original dataset for endothelial cells, fibroblasts, macrophages, and tumor cells in simulated tumor cells. (g) Evaluation metrics benchmarking MisTIC with FastReseg and ResolVI including mean absolute errors for positive transcripts and negative transcripts and AUROC.

For a given transcript initially assigned to a segmented cell, the model evaluates the plausibility of reassignment to a neighboring cell by computing three features: 1. Spatial proximity: A score measuring how close the transcript is to the boundary of the neighboring cell of a different cell type under consideration; 2. Gene expression compatibility: A score reflecting how unlikely it is for the currently assigned cell type to express the transcript’s gene; and 3. Neighborhood expression support: A score capturing how highly the gene is expressed in neighboring cells of a different cell type. These spatial and transcriptomic features then serve as input to the core probabilistic model of MisTIC.

To guide the learning of the model parameters, we use cell group annotations as supervision guidance. Intuitively, correct reassignments should sharpen the gene expression signature of each cell, thereby enhancing the separability of cell groups. Therefore, after transcripts are reassigned, we aggregate gene expression profiles at the cell level and let the updated cell-by-gene matrix pass through a logistic regression classifier to predict the annotated cell groups which are used in the loss function. The cross-entropy between the predicted cell types and the cell type annotations provide signals on how to update all the learnable parameters in the model. To enable joint training, we further cast the problem under a variational Bayes framework by maximizing an evidence lower bound (ELBO) objective (See Methods section for details). The training is accomplished by dividing the tissue into patches and performing stochastic gradient descent (SGD) (21) (Fig. 2b).

After training, the model outputs a reassignment probability for each individual transcript, representing the likelihood that the transcript should be reassigned to a neighboring cell. These reassignment probabilities can be thresholded or used directly in downstream analyses to refine gene expression quantification, improve cell-type classification, and enhance the accuracy of spatial transcriptomic interpretations (Fig. 2c). In addition, the cell level summary of these reassignment probability values can also be used as a measure for the quality of segmentation.

### MisTIC pinpoints misassigned transcripts

To validate MisTIC’s efficacy, we generated a series of simulated datasets based on a publicly available MERSCOPE hepatocarcinoma (HCC) dataset (See Data Availability for detail). The overarching goal of the simulation process is to create synthetic datasets with simulated tumor cells that have both known misassigned (positive) transcripts and correctly assigned (negative) transcripts, while retaining biologically relevant information so that the simulation results can reflect the performance on real datasets as closely as possible.

As tumor cells are densely populated in contiguous regions in the MERSCOPE HCC sample, those in the core region provide a relatively clear background with minimal inherent misassignment transcripts from cells of other types, while tumor cells close to the tumor boundaries are more susceptible to transcript misassignment errors. Based on this observation, we then generated a synthetic dataset featuring an artificial interface between tumor cells and simulated non-tumor cells for the tumor region. The tumor cells in this interface were from the core tumor region, while the simulated non-tumor cells were created by permuting tumor cells originally located within the tumor boundaries and swapping their transcripts with those from non-tumor cells, thereby modifying their expression profiles. This artificially generated tumor:non-tumor interface ensures that the target tumor cells have a low baseline probability of containing misassigned transcripts from non-tumor cells. The simulated tumor cells were then synthesized based on the tumor cells in this interface. To simulate positive transcripts, we injected synthetic transcripts within the vicinity of the boundaries of tumor cells. The identities of those transcripts are sampled from differentially expressed genes from the neighboring cell types. As a negative control, we also simulated tumor-specific transcripts into the same tumor cells with their locations being randomly scattered within target cells. To get a systematic assessment on the performance of different transcript misassignment correction tools, we further varied the maximum number of synthetic transcripts per tumor cell from 5 to 30 with an increment of 5 transcripts, resulting in 6 datasets in total. In Fig. 2d, we show two examples of the resulting simulated tumor cells with added transcripts. (See Methods section for details on synthetic data generation).

We first verified our simulation strategy by visualizing the dataset with 30 added transcripts using UMAP (46, 47) (Fig. 2e). Note that in the leftmost panel, the “Simulated Tumor” cell type indicates the set of tumor cells that will contain added transcripts during the synthetic data generation. By comparing the noisy dataset with the original dataset in Fig. 2e, we see that the two populations of tumor cells overlap fairly well in the original dataset. On the other hand, there is a significant portion of tumor cells with added transcripts that deviate from their intact counterparts, reflecting the success of our simulation strategy.

We then applied MisTIC, ResolVI (24), and FastReseg (25) to this dataset and again visualized the resulting corrected datasets via UMAP (Fig. 2e). Strikingly, MisTIC almost restored the relationship between the two populations of tumor cells back to their original state. Interestingly, from the UMAP embedding, we see that other cell type populations including immune cells and stromal cells appear more separated. This was likely due to the removal of the inherent misassignment errors in the original dataset among those cell types. In sharp contrast, significant deviation of the simulated tumor cells from the other tumor cells can still be observed from the datasets generated by ResolVI and FastReseg with the UMAP embeddings between the noisy dataset and the FastReseg-corrected dataset being almost identical. This suggests an overly conservative correction of FastReseg. It is worth noting that during training, we combined the two populations of tumor cells into one cell type. Therefore, all methods are agnostic of the synthetic data generation process.

We further investigated the degree to which the expression of non-tumor genes in simulated tumor cells decreased following each method. We selected a set of marker genes (See Methods for details) for endothelial cells, fibroblasts, macrophages, as well as tumor cells, computed by their expression levels within tumor cells, and compared them with the expression levels in the original dataset (Fig. 2f). Therefore, for non-tumor genes, a higher ratio would indicate a higher degree of contamination, whereas for tumor genes, the ratio should be around 1. As expected, for non-tumor genes, the expression ratios all exceed 1 for the noise injected dataset, further corroborating the effectiveness of our simulation strategy (one-sided proportion test P value<2.2E-16 for all non-tumor genes). Furthermore, a significant decrease can be observed for both MisTIC-corrected and ResolVI-corrected datasets (one-sided proportion test P value<2.2E-16 for both methods and all non-tumor genes). Aligning with our observation from Fig. 2e, the result from FastReseg is almost identical to the uncorrected dataset (one-sided proportion test P values=0.19, 0.28, and 0.11 for endothelial, fibroblasts, and macrophages, respectively). Reassuringly, none of the methods reduced the expression ratios of tumor-specific genes (one-sided proportion test P values=0.97, 1, and 0.54 for MisTIC, ResolVI, and FastReseg, respectively).

To systematically benchmark the performance of the three methods, we applied them to all six simulated datasets. We first evaluated their performance on the expression level by computing mean absolute error (MAE) (48) (Fig. 2g). Specifically, for a tumor cell with synthetic transcripts, we computed the proportion differences between the expression levels of genes with added transcripts in the corrected dataset and those in the noisy dataset. For simulated transcripts corresponding to non-tumor genes (positive transcripts), the expected proportion difference is 1, as ideally the method should remove all the added transcripts. Conversely, for simulated transcripts corresponding to tumor specific genes (negative transcripts), the expected proportion difference is 0 since these are genes that should not be corrected. Based on this intuition, we then computed MAEs for both positive transcripts and negative transcripts. From Fig. 2g, we can see that MisTIC outperformed both ResolVI and FastReseg by a large margin under both circumstances (one-sided T-test P value<0.01 for all datasets and both methods). Utilizing the positive and negative labels for the synthetic transcripts, we then benchmarked their performance at individual transcript level using Area Under the Receiver Operating Characteristic (AUROC) (48). However, since ResolVI operates only on cellular level, rendering inspecting the results on molecular level almost impossible, we only compared MisTIC with FastReseg. With nearly perfect AUROC, MisTIC surpassed FastReseg, which was unable to detect the synthetic transcripts at an appreciable level.

In addition to testing MisTIC against two existing methods for transcript correction in imaging-based SRT data, we also applied two leading single-cell doublet detection methods, Scrublet (49) and DoubletDetection (50) to the six simulated datasets with various configurations. We expected these methods to have limited sensitivity in detecting low levels of transcript contamination due to the single-cell-based assumption of a 1:1 ratio of transcript contribution between two cells, along with the algorithms’ inability to use spatial information, which we confirmed by testing across a variety of parameters, with all of them resulting in low precision, recall, and F_1_ scores (51) relative to MisTIC (Fig. S3). Specifically, the top performing combination of parameters (selected based on F_1_ score) for Scrublet across the six test datasets have an average precision, recall, and F_1_ score of 0.63, 0.65, and 0.64, respectively. For the top performers of DoubletDetection, the average precision, recall, and F_1_ scores are 0.71, 0.14, and 0.22, respectively. In sharp contrast, the average precision, recall, and F_1_ scores for MisTIC are 1.0, 0.93, and 0.96, respectively.

Together, the simulation study demonstrated the superiority of MisTIC in identifying and correcting misassigned transcripts when compared with algorithms with a similar purpose.

### MisTIC improves cell typing and differential analysis

Analysis of imaging-based SRT data generally starts with unsupervised clustering and annotation of the cell clusters based on reference datasets or marker genes. The contamination from transcript misassignment can confound both processes. In this section, we evaluated MisTIC’s performance in mitigating this problem in a Xenium LUAD dataset (See Data Availability for details) and benchmarked it against ResolVI and FastReseg.

ResolVI was run using the ‘semi-supervised’ mode with cell type annotations, and parameters following recommended values. FastReseg was run with cell type annotation using default parameters in 2D mode with a pixel size of 1. Overall, MisTIC reassigned 3.7% transcripts and removed 8.4% transcripts that cannot be confidently reassigned. FastReseg reassigned and removed 0.03% transcripts in total. Since ResolVI does not correct data at the transcript level, the per-cell change in gene expression relative to library size was calculated. On average, gene expression levels changed by 13% per cell.

We calculated the UMAP embeddings of the SRT data before and after correction. Similar to the results from the HCC MERSCOPE data (Fig. 2e), non-tumor cells in the LUAD Xenium dataset did not show clear separation in the UMAP space (Fig. 3a). In addition, a population of cells resembling both tumor cells and macrophages was observed between the two groups (Fig. 3a).

**Fig. 3.**
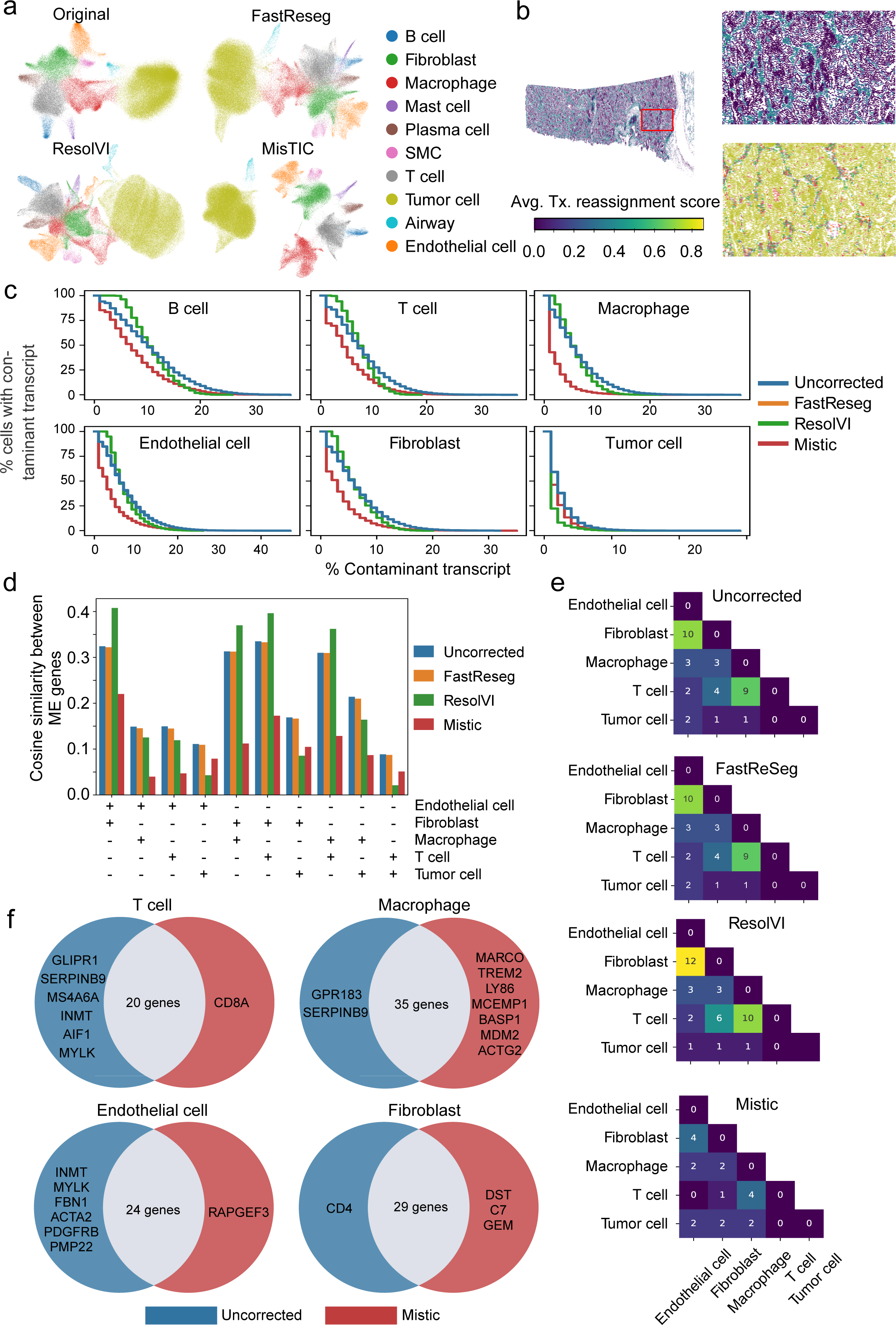
MisTIC enhances cell typing and differential analysis. (a) UMAP embeddings of original and corrected SRT data (FastReseg, ResolVI, MisTIC). (b) Distribution of average transcript reassignment scores across cells, with each cell represented by its centroid. Cells were colored based on the transcript reassignment score and cell type in the upper and lower panel, respectively. (c) Cumulative percentage of contaminant transcripts relative to total transcript counts in different cell types, shown for original and corrected SRT data (FastReseg, ResolVI, MisTIC). (d) Cosine similarity between pairs of cell-type–specific marker gene sets expressed within the same cell. For each gene set pair, average expression was calculated in cells with nonzero expression of at least one marker gene. (e) Heatmap of shared differentially expressed genes (DEGs) between cell types, identified using the Wilcoxon rank-sum test in original and corrected SRT data (FastReseg, ResolVI, MisTIC). DEGs were selected based on log fold-change ≥ 0.6, FDR-adjusted P-value ≤ 0.05, and expression in at least 25% of cells within the corresponding cell type. (f) Venn diagram comparing shared DEGs before and after correction by MisTIC.

This suggests that transcript misassignment-induced contamination confounds cell clustering. Correction using ResolVI or FastReseg did not resolve this issue, as non-tumor cells remained intermixed. In contrast, MisTIC correction effectively separated non-tumor cells and distinguished tumor cells from macrophages (Fig. 3a), indicating that MisTIC improves clustering by identifying and reassigning or removing contaminant transcripts. We then evaluated the spatial distribution of correction intensity by these methods at transcript (MisTIC and FastReseg) or at gene level (ResolVI), calculated as change in gene expression relative to library size in each cell. After MisTIC correction, cells in regions with tumor-stromal or tumor-immune interfaces (Fig S1a, Fig. 3b) had higher correction intensity compared to other regions. This pattern was not present in data corrected by FastReseg or ResolVI (Fig. S4 a and b).

Next, we evaluated the correction effectiveness of these methods in various cell types, including immune, stromal, and tumor cells. Similar to the analysis in Fig. 1c, we calculated the total counts from contaminating genes in cells and evaluated the cumulative distribution of cells with various levels of contamination. Compared to uncorrected data, FastReseg did not cause any discernible changes in the distribution of contaminant transcripts. ResolVI reduced the number of cells with >10% contamination but increased the number with low-level contamination. This is likely because the correction method failed to detect erroneous gene counts when the errors were too small to cause substantial transcriptional changes, or because the correction was incomplete, reducing but not fully eliminating the expression of misassigned transcripts. MisTIC outperformed ResolVI in all cell types except tumor cells, where contamination was low and ResolVI performed slightly better. MisTIC reduced the number of contaminated cells across all contamination levels, doubled the number of cells without contamination, and led to an over 50% reduction in the numbers of cells with contamination levels at 10% or above, in all cell types except tumor cells (Fig. 3c).

We also evaluated each method’s performance in reducing cross-contamination between cell type pairs, measured as the cosine similarity between two mutually exclusive gene sets expressed in the same cell (24) (See Methods for details). Compared with the uncorrected data, FastReseg correction only slightly reduced the cross-contamination between cell types. ResolVI was effective in 6 out of 10 cell type pairs but increased the cross-contamination in 4 cell pairs involving endothelial cells, fibroblasts, macrophages or T cells. In contrast, MisTIC consistently reduced cross-contamination in all cell pairs and was only slightly out-performed by ResolVI in cell pairs involving tumor cells (Fig. 3d).

Lastly, we compared the differential expression analysis results from data corrected by each method. In the uncorrected data, we identified substantial overlap in differentially expressed genes (DEGs) between fibroblasts and endothelial cells, as well as between macrophages and T cells. Corrections by FastReseg and ResolVI did not reduce the number of shared DEGs. In contrast, MisTIC correction reduced DEG overlap in both cell type pairs (Fig. 3e). To distinguish DEGs arising from transcript contamination versus those reflecting genuine shared biology, we referenced LUAD scRNA-seq datasets (52–55) to assess the expression of these DEGs in their corresponding cell types. For example, DEGs shared by fibroblasts and endothelial cells in the uncorrected data included genes rarely expressed in endothelial cells, such as *MYLK*, *FBN1*, *ACTA2*, *PDGFRB*, and *PMP22* (Fig. S5a, Fig. 3f). MisTIC correction removed these fibroblast-specific genes from the endothelial DEG list while preserving shared genes such as *CAVIN1*, *TCF4*, and *ADAMTS1*. Similarly, in the comparison between macrophages and T cells, MisTIC reduced ambiguity by removing AIF1, *CD68*, and *MS4A6A* from T cells, and *SERPINB9* and *GPR183* from macrophages (Fig. S5b, Fig. 3f). Correction by MisTIC also allowed identification of additional DEG from these cell types, compared to uncorrected data, such as *CD8* in T cells, and *MARCO, TREM2,* and *LY86* in macrophages (Fig. 3f).

Taken together, these results showed MisTIC effectively corrected transcript misassignment in imaging-based SRT data and improved cell clustering. In addition, it decreased spurious DEG overlap between related cell types while preserving biologically relevant expression patterns.

### MisTIC enhances cell-cell communication detection

Accurate inference of cell-cell communication (CCC) from imaging-based SRT data critically depends on the fidelity of cell-type-specific expression profiles. These profiles form the basis for quantifying ligand-receptor (L-R) interactions between spatially defined cell types, whether through contact-based CCC, in which ligands and receptors engage directly across adjacent plasma membranes, or paracrine CCC, in which soluble ligands diffuse through the extracellular space to reach nearby receptor-bearing cells (Fig. 4a) (56). Transcript misassignment, a common consequence of segmentation errors, can redistribute transcripts located near cell boundaries to incorrect cells. Such redistribution artificially blurs the molecular distinction between neighboring cells, dilutes genuine L-R signals, and may generate spurious interactions. This problem is especially acute for L-R pairs whose expression is spatially localized near intercellular boundaries, where even modest errors can substantially alter the inferred interaction network.

**Fig. 4.**
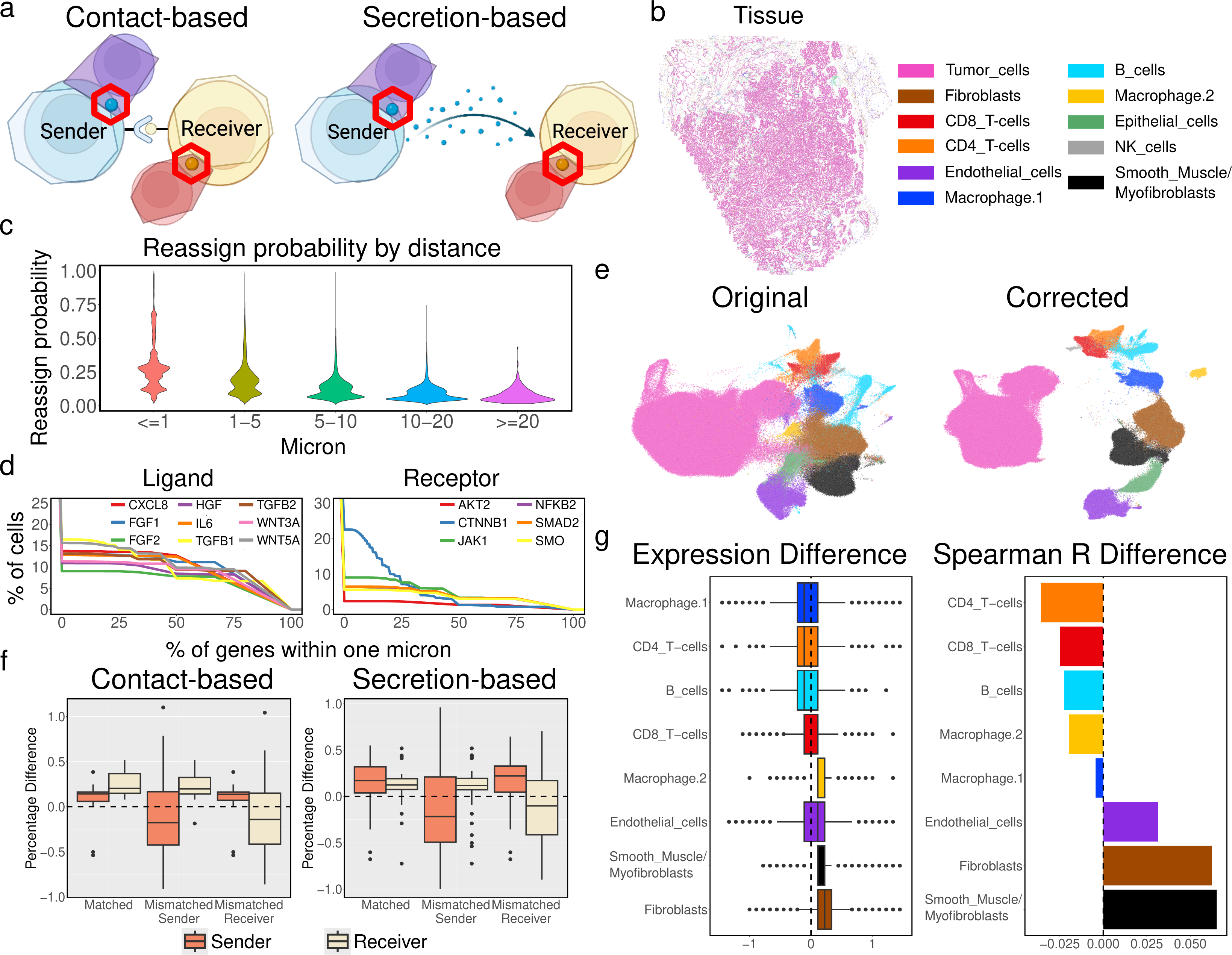
MisTIC enhances cell-cell communication detection. (a) Illustration of contact-based CCC and secretion-based CCC. (b) MERSCOPE prostate cancer SRT data with cell type annotation. (c) Reassignment probability for each EMT-related transcript versus its distance to its nearest neighbor cell. (d) Cumulative percentage of cells that contain boundary-proximal EMT-related transcripts. (e) UMAP embeddings of the original and MisTIC-corrected SRT data. (f) Percentage changes in expression profiles by sender-receiver type with two negative control groups by contact-based CCC and secretion-based CCC. (g) Overall expression changes and changes in Spearman correlations between ligand expression in non-tumor cells and receptor expression in tumor cells.

MisTIC addresses this challenge by probabilistically reassigning boundary-proximal transcripts to their most likely cell of origin based on spatial proximity and local transcript context. By restoring sharp expression boundaries and more coherent spatial patterns, MisTIC is expected to strengthen true CCC signals while suppressing false positives arising from misassignment. To evaluate this hypothesis, we analyzed a publicly available MERSCOPE dataset of human prostate cancer (See Data Availability for detail) (Fig. 4b). As an illustrative example, we focused on epithelial-mesenchymal transitions (EMT) in tumor cells, a process central to cancer progression and metastasis that is often modulated by paracrine (57) and juxtacrine (58) communication with the tumor microenvironment. EMT can be induced by a variety of immune-and stroma-derived ligands acting upon corresponding receptors in tumor epithelial cells to trigger transcriptional reprogramming toward a mesenchymal phenotype (30). We identified a set of well-known ligands expressed by immune and stromal populations with known EMT-inducing activity including *WNT5A*, *WNT3A*, *TGFB1*, *TGFB2*, *FGF2*, *IL6*, *CXCL8*, *HGF*, and *FGF1* (59–66) as well as EMT markers or drivers in tumor cells including *JAK1*, *AKT2*, *SMO*, *CTNNB1*, *SMAD2*, and *NFKB2* (*61–69*). We first examined how the reassign probabilities for those genes computed by MisTIC vary with the distance from the nearest cell boundary (Fig. 4c). As expected, MisTIC assigns higher probabilities to boundary-proximal transcripts while for transcripts that are further away the probabilities are close to 0. For each gene, we then further quantified the fraction of transcripts located within 1 μm of the nearest cell boundary where most of the correction occurs (Fig. 4d). We found that approximately 10-15% of EMT-related transcripts in both ligand-expressing and receptor-expressing cells fell within this boundary-proximal zone, suggesting that a substantial portion of EMT-relevant signals are at risk of being distorted by segmentation errors.

We next used CellChat v2 (70) which extends CellChat (71) by incorporating spatial information to infer candidate sender-ligand:receiver-receptor quadruples and classified them into contact-based or secretion-based interactions. For each quadruple, we assigned ligand and receptor genes to a cell type based on their expression, and we computed ligand and receptor expression as the sum of transcripts within the respective sender and receiver cell types. Applying MisTIC to the same dataset yielded corrected expression values, enabling direct calculation of percentage changes between the original and corrected data. A quick visual inspection comparing UMAPs before and after correction revealed crisper intercellular borders, suggesting reduced “bleed-through” of transcripts between adjacent populations (Fig. 4e). To evaluate if MisTic correction was enriched in true CCC quadruples , we generated two negative control groups: a “mismatched sender” group in which sender cell types in each quadruple were permuted to cell types not expressing the ligand, and a “mismatched receiver” group in which receiver cell types in each quadruple were permuted to cell types not expressing the receptor. Quadruples in these negative groups represent false positives in CCC predictions. If boundary-driven false positive expression is present, we expect a reduction after MisTic correction.

To further emphasize the expected directional changes, we aggregated the percentage changes in expression profiles by sender-receiver type (Fig. 4f). In the matched, biologically relevant quadruples, we observed consistent increases in ligand and/or receptor expression after MisTIC correction, reflecting sharper cell-type-specific expression boundaries (P-values: 0.039, 7.6e-6 in ligands and receptors in contact-based CCCs, respectively and 0.0005, 0.0004 in ligands and receptors in secretion-based CCCs). In contrast, both negative control groups showed decreases in expression (P-values: 0.027 and 0.004 in mismatched legends and receptor in contact-based CCCs, respectively and 1.8e-6 and 0.0007 in mismatched legends and receptor in secretion-based CCCs), consistent with the removal of spurious boundary-derived signals. These effects were evident in both contact-based and secretion-based CCCs. These observations further corroborate that MisTIC selectively reinforces biologically plausible CCCs while suppressing artifacts. Importantly, application of the same procedure to an independent CosMX dataset of human pancreas tissue (See Data Availability for details) produced qualitatively similar patterns, indicating that the observed effects are not dataset-specific (Fig. S6).

Given that our initial motivation was to use EMT as an illustrative case of how segmentation errors can obscure biologically important CCCs, we next revisited EMT-related ligand-receptor interactions in detail. EMT is not only a hallmark of cancer progression but also a process that integrates signals from multiple microenvironmental sources, including direct contact with stromal cells and paracrine stimulation from immune populations (59–61, 64, 65). Because these signals operate at different spatial scales and degrees of cell-type specificity, we reasoned that evaluating MisTIC’s impact at two complementary levels would provide a more complete picture of how correction influences CCC readouts.

At the cell-type level, we sought to capture broader trends in ligand production across stromal and immune populations. This perspective reflects the idea that many EMT-inducing signals are regulated at the population scale and may exert cumulative effects on tumor cells over both short and intermediate distances. By quantifying the expression of EMT-related ligands in each immune or stromal cell type before and after correction, we could assess whether MisTIC systematically sharpened these population-level signatures. At the neighbor-level, we aimed to detect more localized and spatially coordinated interactions, which are especially relevant for juxtacrine signaling and for paracrine signals with limited diffusion. Here, for each tumor-immune/stromal neighbor pair, we computed the Spearman correlation between ligand expression in the non-tumor cell and receptor expression in the tumor cell. This measure reflects the degree to which cells in local microenvironments are functionally coupled in their expression. In the presence of misassignment errors resulting in mixing transcripts across boundaries, this coupling can be weakened.

Post-correction, fibroblasts, smooth muscle cells/myofibroblasts, and endothelial cells exhibited consistent increases in both cell-type level ligand expression and neighbor-level ligand-receptor correlation, in sharp contrast with myeloid cells and lymphoid cells. Given their well-established roles in promoting EMT (59, 62, 68, 69, 72), this observation supports the claim that MisTIC enhances both broad, population-scale signaling capacity and fine-scale, spatially coherent interactions. By correcting transcript misassignment, MisTIC effectively sharpens the attribution of EMT-inducing ligands to the most biologically plausible cellular sources while reducing spurious contributions from unrelated immune populations.

### MisTIC facilitates RNA localization analysis

One unique advantage of imaging-based SRT technologies is their ability to capture nuclear and cytoplasmic compartment information for each cell. This enables the analysis of transcript distribution between these compartments, offering insights that complement traditional average-based gene expression analyses. Specifically, it could shed light on transcript regulation dynamics related to mRNA transport, retention, and degradation. Misassignment of transcripts can distort cytoplasmic distribution, much like it introduces errors in cell typing, differential expression analysis, and predictions of cell-cell interactions. In this section, we evaluated MisTIC’s performance in reducing uncertainty in transcript localization using benchmark datasets. Notably, we found that cancer-associated fibroblasts were linked to cytoplasmic depletion of genes involved in T cell activation within T cells.

The earlier Xenium platform was limited to profiling only a few hundred genes, making unbiased assessments of transcript distribution and subsequent benchmarking challenging. To address this, we expanded our analysis to the Xenium Prime platform, which profiles 5,000 genes. Two datasets were used: one from primary dermal melanoma and another from prostate adenocarcinoma (See Data Availability for details). Due to the sparsity of SRT data at the per-gene level in Xenium Prime, we calculated the cytoplasmic fraction of each gene at the dataset level. This was defined as the ratio of transcripts located in the cytoplasmic region to the total number of transcripts within cells, considering only high-quality cells. MisTIC correction did not substantially alter the overall cytoplasmic fraction of genes compared to uncorrected data. Among the 5,000 genes, only 440 showed a deviation of at least 5% in cytoplasmic fraction following MisTIC correction (Fig. 5a).

**Fig. 5.**
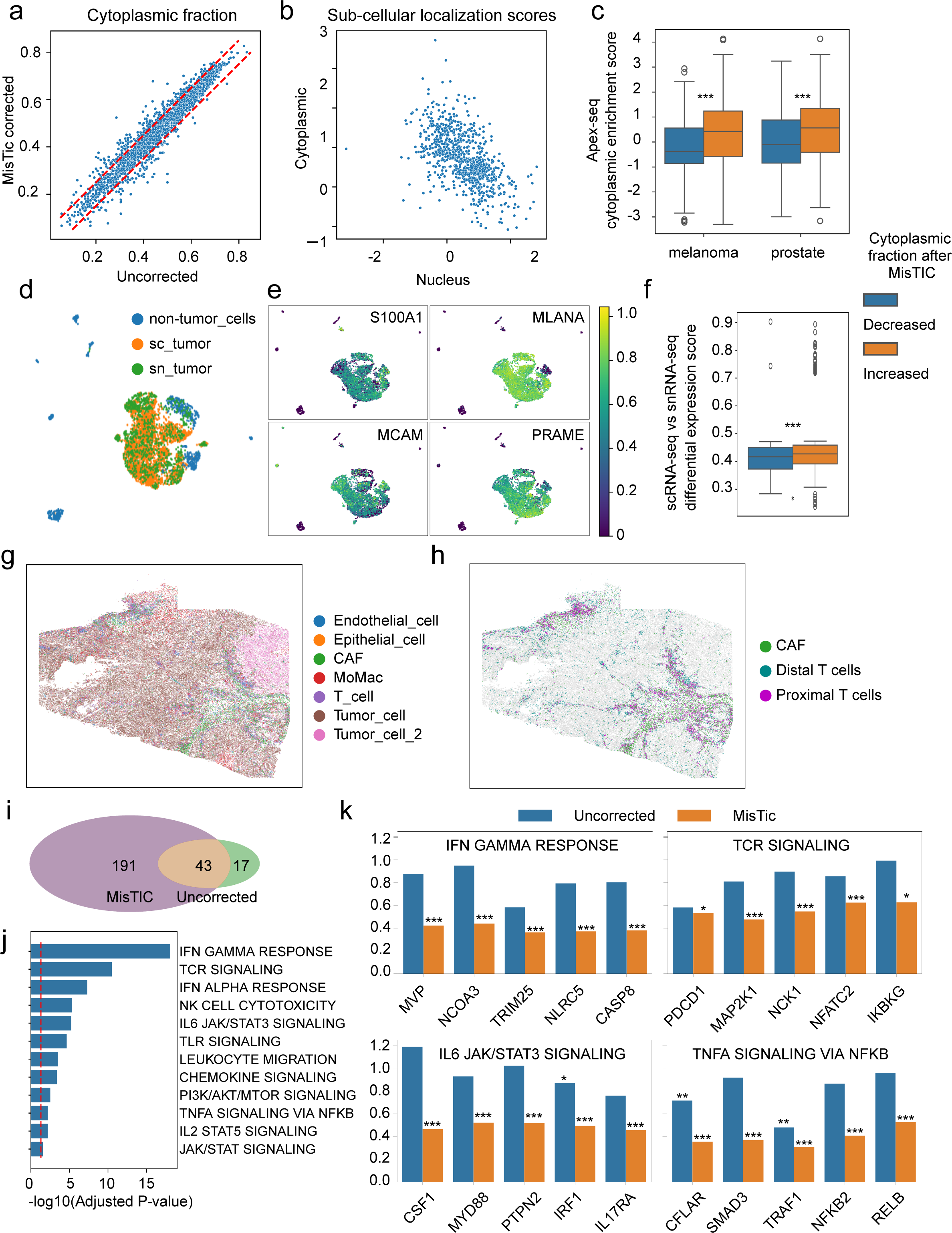
MisTIC facilitates RNA localization analysis. (a) Scatter plot of cytoplasmic fractions of genes in original and MisTIC-corrected Xenium Prime data from human melanoma tissues. (b) Scatter plot of cytoplasmic and nuclear localization scores for 875 Xenium Prime genes included in the APEX-seq reference dataset. (c) Comparison of APEX-seq–derived cytoplasmic enrichment scores in transcripts with increased (orange) or decreased (blue) cytoplasmic fractions after MisTIC correction. Cytoplasmic enrichment scores were calculated as the difference between cytoplasmic and nuclear localization scores. Significance was estimated using Student’s t-test (*P ≤ 0.05; **P ≤ 0.01; ***P ≤ 0.001). (d) UMAP embedding of reference sc/snRNA-seq data from human melanoma tissues, integrated using Harmony. (e) Expression of key melanoma marker genes detected in both SRT and sc/snRNA-seq data. Expression values were scaled from 0 to 1. (f) Comparison of differential expression scores in transcripts with increased (orange) or decreased (blue) cytoplasmic fractions after MisTIC correction. Scores were calculated by comparing tumor cells from scRNA-seq with those from snRNA-seq. Significance was estimated using Student’s t-test (*P ≤ 0.05; **P ≤ 0.01; ***P ≤ 0.001). (g, h) Spatial distribution of all cell types (g) and CAFs with proximal/distal T cells (h) in the melanoma SRT dataset. (i) Venn diagram of T cell transcripts with significantly decreased cytoplasmic presence in proximal versus distal T cells, before and after MisTIC correction. Significance was estimated using Fisher’s exact test (P ≤ 0.05). (j) Odds ratios for cytoplasmic presence of representative genes in pathways related to T cell function or inflammatory responses, comparing proximal versus distal T cells. Values from original (blue) and MisTIC-corrected (orange) data are shown. Significance was estimated using Fisher’s exact test (*P ≤ 0.05; **P ≤ 0.01; ***P ≤ 0.001). (k) Enrichment analysis of 191 genes uniquely identified by MisTIC.

To further evaluate MisTIC’s performance, we used cytoplasmic localization data from an APEX-seq dataset (73). APEX-seq is a proximity-based RNA sequencing method that uses the peroxidase enzyme APEX2 to label RNA near specific subcellular locations, enabling simultaneous measurement of gene expression and subcellular localization. In this dataset, 3,289 genes were enriched in at least one of the 9 assayed subcellular locations, including nucleus and cytoplasm, and 875 of these were present in the SRT data. Cytoplasmic and nuclear localization scores of these genes, calculated from the location enrichment fold changes, were negatively correlated (Fig. 5b) in the APEX-seq data. Then, to assess MisTIC’s response in the context of APEX-seq data, we categorized the 875 genes into two groups: those with increased cytoplasmic fraction after correction and those without. In both the melanoma and prostate datasets, genes with increased cytoplasmic fraction post-MisTIC correction also had significantly higher cytoplasmic localization scores in APEX-seq (P-values: 1.14e-9 and 5.58e-5, respectively; Fig. 5c). These results showed that MisTIC correction led to improvements in RNA localization when tested against reference datasets.

In another benchmarking test, we used matched scRNA-seq and single-nucleus (sn)RNA-seq data as a reference to assess MisTIC’s correction performance. This approach is based on the rationale that snRNA-seq typically yields fewer reads for genes enriched in the cytoplasm compared to scRNA-seq. Therefore, differential expression analysis between scRNA-seq and snRNA-seq data from the same cell type can reveal genes with a tendency for cytoplasmic enrichment. We analyzed a dataset in which a melanoma tumor sample was split for parallel scRNA-seq and snRNA-seq profiling (74). Tumor cells from both modalities were highly similar, and key melanoma marker genes—including *S100A1*, *MLANA*, *MCAM*, and *PRAME*—were consistently highly expressed (Fig. 5d-e). Differential expression analysis comparing tumor cells from scRNA-seq and snRNA-seq identified 4,756 genes, of which 661 were also expressed in tumor cells from the SRT dataset. Among these, genes that showed increased cytoplasmic fraction following MisTIC correction also exhibited higher differential expression scores, indicating elevated expression in the scRNA-seq data (P-value = 2.79e-6) (Fig. 5f). Together with the validation using the APEX-seq data, this result suggests that MisTIC correction enhances the robustness of gene localization predictions derived from SRT data.

To demonstrate the transcript subcellular localization analysis powered by MisTIC could drive new biological findings, we evaluated the link between cancer-associated fibroblasts (CAFs) and T cell function in melanoma. We focused on regions with abundant tumor cells from the melanoma Xenium Prime dataset (Fig. 5g) and categorized T cells into two groups: those located within 10 µm of at least two CAFs (Proximal T cells), and those without nearby CAFs (Distal T cells) (Fig. 5h). For each gene, we calculated the odds ratio (OR) comparing the likelihood of transcripts being located in the cytoplasmic region versus the nuclear region between these two T cell groups. A lower OR indicates that the transcript is depleted from the cytoplasm of T cells near CAFs. We performed this analysis both before and after MisTIC correction. In the uncorrected data, 60 genes had OR values less than 1, while in the MisTIC-corrected data, this number was 234 (Fig. 5i). The genes uniquely identified in the corrected data were enriched in pathways related to T cell activation, interferon responses, and inflammation (Fig. 5j). For many of these genes, OR values were significantly reduced following MisTIC correction, sometimes by more than 50% (Fig. 5k). Upon closer inspection, we confirmed that this reduction was primarily due to the correction of transcripts corresponding to T cell genes to T cells distant from CAFs (∼352 thousands corrected transcripts in distal T vs ∼184 thousands in proximal T). Notably, most of these genes were not detected through differential expression analysis comparing the two T cell groups. Importantly, among these pathways, only interferon responses and IL2 signaling were identified via differential expression analysis comparing proximal with distal T cells. These findings suggest that, beyond repressing gene expression, CAFs may suppress T cell function by reducing the cytoplasmic presence of mRNA. This, in turn, could contribute to diminished inflammatory and cytotoxic activity in T cells. Importantly, neither differential analysis on gene expression nor subcellular localization analysis without MisTIC correction could identify these genes.

## Discussion

Cell segmentation errors can have a significant impact on downstream analyses of imaging-based SRT data and limit the usability of such powerful platforms. Cells with misassigned transcripts conceptually resemble ‘cell doublets’ in the scRNA-seq term but are fundamentally different for two reasons: 1. ‘Cell doublets’ are created by two (or more) cells being processed under one barcode, and thus the gene expression could deviate greatly from any of the component cells. In SRT data, expression profiles of cells with misassigned transcripts are much less impacted, as the numbers of contaminating transcripts are relatively small; 2, The number of ‘cell doublets’ is usually very small (∼1-15%, (75–77)) in a single-cell/nuclei dataset, however in SRT, the number of cells affected by cell segmentation error can be much bigger depending on the cell segmentation and the tissue type. For these reasons, single-cell based cell doublet detection algorithms assuming a ‘cell doublet’ are rare and a result of 1:1 mix of two cell type profiles are quite likely to perform poorly in SRT data. To our knowledge, there are two methods for cell segmentation/transcript assignment error correction. ResolVI employs a variational autoencoder to decompose each cell’s expression profile into intrinsic signal, neighbor spillover, and background contamination. While powerful, this formulation operates at the gene-expression level rather than at the transcript level, making it inherently insensitive to the spatial granularity of individual molecules and prone to global over-smoothing. FastReseg, in contrast, attempts transcript-level reassignment through a multistep workflow built around log-likelihood ratios; however, its heavy reliance on ad hoc heuristics and its evaluation on a single CosMX dataset raise concerns about robustness and generalizability. In this study, we introduce MisTIC, a fundamentally different and conceptually unified probabilistic framework that models transcript misassignment directly at the transcript level. Rather than relying on post hoc smoothing or heuristic filtering, MisTIC explicitly represents the spatial and expression-derived uncertainties governing transcript-to-cell assignment. This principled formulation resolves a major, yet under-addressed, bottleneck in imaging-based SRT analysis, providing a scalable and generalizable foundation for improving data fidelity and downstream biological interpretation.

The core innovation of MisTIC lies in its formulation of the transcript reassignment problem within the framework of the variational Bayesian method, where the misassignment probability of each detected transcript is modeled as the output of the variational posterior distribution. By carefully crafting and incorporating three biologically-informed features into the model— spatial proximity, gene expression compatibility, and neighborhood expression support—MisTIC can identify transcripts with high misassignment probability while preserving confident assignments. Furthermore, it can scale up to handle the massive datasets generated by imaging-based SRT platforms, which often contain tens of millions to hundreds of millions of detected transcripts. For example, one epoch of the training process on the MERSCOPE dataset which contains more than 180 million detected transcripts takes about 55 minutes on a CPU. Inference of the transcript reassignment probability for all detected transcripts once the training is done takes approximately 1 minute. The peak memory usage is less than 200GB. This computational efficiency ensures that MisTIC can be routinely applied to large-scale studies without prohibitive computational costs. Additionally, the method’s design allows for straightforward integration into existing analysis pipelines, minimizing the barrier to adoption by the broader research community.

Through our comprehensive validation across multiple datasets and tissue types, we showed that transcript misassignment is a pervasive issue in imaging-based SRT data, with significant implications for biological interpretation. We further demonstrated that MisTIC, via substantially improving the accuracy of transcript-to-cell assignments, can lead to more reliable downstream analyses and biological insights. Specifically, MisTIC enables a more accurate characterization of cellular heterogeneity, enhanced resolution of spatial gene expression patterns, improved cell-cell communication prediction, and uncovered key insights into the mRNA sublocalization in T cells in response to CAF that are masked by transcript misassignment in the uncorrected data.

While MisTIC shows robust performance across diverse datasets, several considerations should guide its application. Since the method does not touch upon the predicted cell polygons, its effectiveness depends on the quality of the initial delineation of cell boundaries. For coarse cell boundaries, we envision multiple runs of MisTIC where the transcript assignments as well as cell type annotations are iteratively refined might alleviate the issue. As the field advances from single z-level 2D segmentation toward multilevel or fully 3D cell segmentation, MisTIC could be further extended to take advantage of these developments. In such a setting, the framework could be adapted to jointly refine 3D cell contours, transcript assignment in 3D space. Integration of morphological features, tissue architecture constraints, or prior biological knowledge about gene expression patterns could also potentially enhance the model’s ability to distinguish correct from incorrect assignments.

As imaging-based SRT technologies continue to evolve toward higher resolution and throughput, robust computational methods for data quality control will become increasingly important. MisTIC provides a foundation for addressing one critical aspect of data quality, and similar probabilistic approaches could be developed to address other systematic errors in spatial omics data. The open-source availability of MisTIC facilitates community adoption and further development, supporting the continued advancement of spatial transcriptomics analysis methods. With continued advances in multimodal cell surface staining and cell segmentation algorithms, segmentation accuracy is anticipated to improve to the extent that post-hoc correction may no longer be required. Under such conditions, MisTIC could be run post segmentation and generate quality control metrics that quantify segmentation accuracy at the single-cell level through transcript reassignment scores.

In conclusion, MisTIC represents a significant step forward in addressing a fundamental challenge in imaging-based SRT data analysis. By providing an effective, scalable, and user-friendly solution for transcript misassignment correction, MisTIC enhances the reliability and biological interpretability of spatial transcriptomics studies, ultimately supporting more accurate scientific discoveries from these powerful experimental platforms.

## Methods

### Technical overview of MisTIC

MisTIC is a Bayesian hierarchical model that frames the transcript misassignment correction problem as a sequence of binary classification problems. For each detected transcript, MisTIC computes a probability on whether or not it should be reassigned to its neighboring cell of a different cell type than the cell it is currently assigned to. Three biologically inspired factors are considered during the computation: 1. Spatial proximity: A score measuring how close the transcript is to the boundary of the neighboring cell of a different cell type under consideration; 2. Gene expression compatibility: A score reflecting how unlikely it is for the currently assigned cell type to express the transcript’s gene; and 3. Neighborhood expression support: A score capturing how highly the gene is expressed in neighboring cells of a different cell type. MisTIC is tailored to the analysis of imaging-based ST datasets (such as 10X Xenium, MERSCOPE, and CoxMX) where the spatial locations of each detected transcript is available. Therefore, MisTIC is not expected to work with data from sequencing-based ST platforms such as Visium HD where the spatial information of transcripts is only available at the spot level. Users are expected to provide cell masks defined via segmentation techniques. Cell type can usually be determined from the aggregated single-cell level expression profiles or from morphological features. The details of the probabilistic model of MisTIC are provided in File S1.

### Synthetic data generation

To ensure that the simulated data reflects biologically relevant transcript distributions across different cell types, we modified an existing MERSCOPE hepatocarcinoma (HCC) scSRT dataset to generate a synthetic dataset. The objective was to simulate transcripts in two categories: a positive category representing missegmented transcripts, and a negative category representing transcripts native to the host cell type.

We selected tumor cells as the host cell type because they densely populated a contiguous region in the liver. To minimize potential confounding from existing misassigned transcripts, only tumor cells located at least 2 μm away from any non-tumor cells (including immune and stromal cells) were used. For tumor cells within 2 μm of non-tumor cells, we randomly reassigned their cell type annotations to non-tumor types. The gene expression profiles of these reassigned cells were then replaced with synthetic values sampled from the DEGs of the corresponding non-tumor cell types. This created an artificial tumor-non-tumor interface, where the host tumor cells have a low baseline probability of containing heterotypic misassigned transcripts.

To simulate misassigned transcripts, we randomly added synthetic transcripts to near-mask regions (defined as transcript-to-mask distance ≤ 1 μm) within each host cell. The gene identity of each added transcript was sampled from the top differentially expressed genes in the nearest non-tumor neighbor. These were labeled as the positive category. As a control, we also added transcripts corresponding to tumor-specific genes to the same host cells, representing the negative category. Multiple synthetic datasets were generated with varying parameters controlling the maximum number of added transcripts per host cell.

### Standard data analysis

Data preprocessing. Quality control, filtering, normalization and cell type annotation was done using standard Scanpy functions. Briefly, we removed cells with fewer than 20 total transcripts from the analysis, and normalized gene expression data using library size normalization and log1p transformation. Unsupervised clustering was done using the Leiden algorithm following dimension reduction by Principal Component Analysis (PCA). Cell type annotation was performed by referencing known cell type markers and single cell atlas portals such as CellxGene (https://cellxgene.cziscience.com/datasets) and the Broad Single Cell Portal (https://singlecell.broadinstitute.org/single_cell). Visualization of cell typing results was performed using the Uniform Manifold Approximation and Projection algorithm (UMAP).

### Marker genes

Marker genes were selected based on literature and differential analysis, and listed below. Endothelial cells: *CLDN5*, *RAMP2*, *VWF*, *EGFL7*, *CLEC14A*, *ACKR1*, *ECSCR*, *CD93*, *PLVA*, *CDH5*; Fibroblasts: *PTGDS*, *FBLN1*, *MMP2*, *SFRP2*, *ACTA2*, *PCOLCE*, *THY1*, *PLAC9*, *COL6A3*, *ELN*; Macrophages: *AIF1*, *CD68*, *FCGR3A*, *MS4A6A*, *MARCO*, *SPI1*, *CD14*, *TREM2*, *MSR1*, *MRC1*, *CD163*, *CSF1R*; Tumor cells: *KRT7*, *SFTA2*, *AGR3*, *EPCAM*, *GPRC5A*, MMP7, *CDH1*, *MET*, *ERBB3*, *LAMB3*.

### Assessment of transcript misassignment in cells

We assessed contamination in gene expression by quantifying the expression of genes not specific to a given cell type. Candidate marker genes were identified from reference scRNA-seq data of human LUAD tissues by filtering differentially expressed genes (DEGs). For each evaluated cell type (B cells, T cells, macrophages, tumor epithelial cells, and fibroblasts), the top five DEGs expressed in fewer than 5% of other cell types were selected as cell-type-specific markers (Shown in Fig. 1a). For each cell, contamination from other cell types was estimated as the ratio of the summed expression of 20 non-specific marker genes (derived from the four non-matching cell types) to the total library size. Similarly, for each potential contaminating cell type, the same measure was calculated using the five marker genes specific to that cell type. We also assessed the contamination using library normalized data, as previously reported (24). We calculated the average expression of cell type-specific marker genes within each cell. Then we computed the cosine similarity between their average expression profiles, to evaluate co-expression between marker sets from two different cell types.

### Cytoplasmic localization score calculation in reference datasets

RNA localization data was extracted from Table S3 from the original Apex-seq paper published in 2019 (73), which contains enrichment log fold-changes (logFC) of a RNA to each assayed subcellular locations, including nucleolus, nucleus, nuclear lamina, nuclear pore, ER lumen, ER membrane, cytosol, mitochondria matrix (MM), and outer mitochondria membrane (OMM). Nuclear pore, ER lumen and ER membrane were removed from consideration due to these structures being close to both nucleus and cytoplasm. To consolidate these scores into a single cytoplasmic score, we calculated the difference between medians of logFC corresponding to cytoplasmic locations (cytosol, MM, OMM) and those to nucleus locations (nucleolus, nucleus, nuclear lamina,). A large value would indicate the RNA’s higher tendency of being enriched in the cytoplasm.

For the reference dataset involving sc/sn RNA-seq of human melanoma tissues, we download processed gene expression matrices containing the sample of interest (74). Data preprocessing and identification of tumor cells was done as previously described in the “Standard data analysis” section. Differential analysis of tumor cells from between snRNA-seq and scRNA-seq was performed using the Wilcoxon Rank Sum test. We followed the original author’s suggestion in their publication regarding the removal of stress related genes from the DEG. In addition, we used a more stringent cutoff for logFC to focus on genes that have the highest difference between the two dataset. The differential expression scores calculated by Scanpy were used as the cytoplasmic localization score in our analysis. A large value would indicate the RNA’s higher tendency of being enriched in the cytoplasm.

### Benchmark study

#### ResolVI

We installed, trained, and performed corrections following the tutorial provided by the authors: https://docs.scvi-tools.org/en/1.3.0/tutorials/notebooks/spatial/resolVI_tutorial.html.

Semi-supervised mode is used whenever cell type annotation is available. Default parameters were used for all analyses.

#### FastReseg

The simulated data were tested using FastReseg to quantify the tool’s ability to detect and correct for missegmented transcripts. In accordance with the author’s recommendations, we focused on varying the SVM score cutoff (-2.3, -2.2, -2.1, -2, -1.9, -1.8, and -1.7), gamma (0.2, 0.3, and 0.4), and cost (1, 10, and 100) parameters, as they are the most likely factors to affect tool performance. An appropriate pixel size for the data was selected and FastReseg was run in 2D-mode. All other parameters were left at default values.

#### DoubletDetection

The six simulated data sets were tested with DoubletDetection v4.3.0 using default values as well as a wide range for the voter threshold (0.1-0.9, by 0.1) parameter and all three available clustering algorithms (louvein, phenograph, and leiden). True positive cells were defined as those tumor cells containing simulated heterotypic transcripts that were accurately detected by the tool out of all tumor cells containing simulated transcripts (positives). False positives were defined as tumor cells detected by the tool as being doublets that did not contain simulated transcripts.

#### Scrublet

Similarly, the six test data sets were processed with Scrublet v0.2.2 using default values as well as varying the threshold parameter (0.1-0.9, by 0.1) and expected doublet rate (0.02-0.2, by 0.02). True positive and false positive populations were defined in the same manner as for DoubletDetection.

### Statistics and Reproducibility

Computations were performed in R (4.4.2) and Python (3.9 and 3.10) programming languages. All statistical tests were two-sided unless otherwise described. For all box plots, box boundaries represent interquartile ranges, whiskers extend to the most extreme data point, which is no more than 1.5 times the interquartile range, and the line in the middle of the box represents the median.

## Data Availability

Datasets. The imaging-based SRT datasets analyzed in this manuscript were obtained from the publicly accessible data portals of 10x Genomics, Vizgen, and NanoString. For each dataset, cell segmentation was carried out by the respective platform using multimodal cell surface staining. A full list of the datasets used in this manuscript can be found in Table S1.

## Code Availability

The MisTIC software is available at https://github.com/yunguan-wang/MisTIC-Wanglab. Detailed instructions on installation and misassignment correction are also provided. The code and data to reproduce the results presented in the manuscript can be accessed via https://doi.org/10.5281/zenodo.16876645.

## Acknowledgements

This project was supported in part by NIH P30 DK078392 Gene Analysis Core of the Digestive Diseases Research Core Center in Cincinnati. CpG Grant, and NIH 1R01GM160515-01/XW & YY.

## Author contributions

Conceptualization: YY, YW, and ED. Data curation: YY, YW, ED, DA, and XX. Formal analysis: YY, YW, ED, DA, and XX. Funding acquisition: YW and ED. Investigation: YY, YW, and ED. Methodology: YY, YW, and ED. Project administration: YY, YW, and ED. Resources: YY, YW, and ED. Software: YY, YW, and ED. Supervision: YY, YW, and ED. Validation: YY, YW, ED, and DA. Visualization: YY, YW, ED, and DA. Writing – original draft: all authors wrote the manuscript.

## Declaration of interests

The authors declare no competing interests.

---

**Fig. S1.**
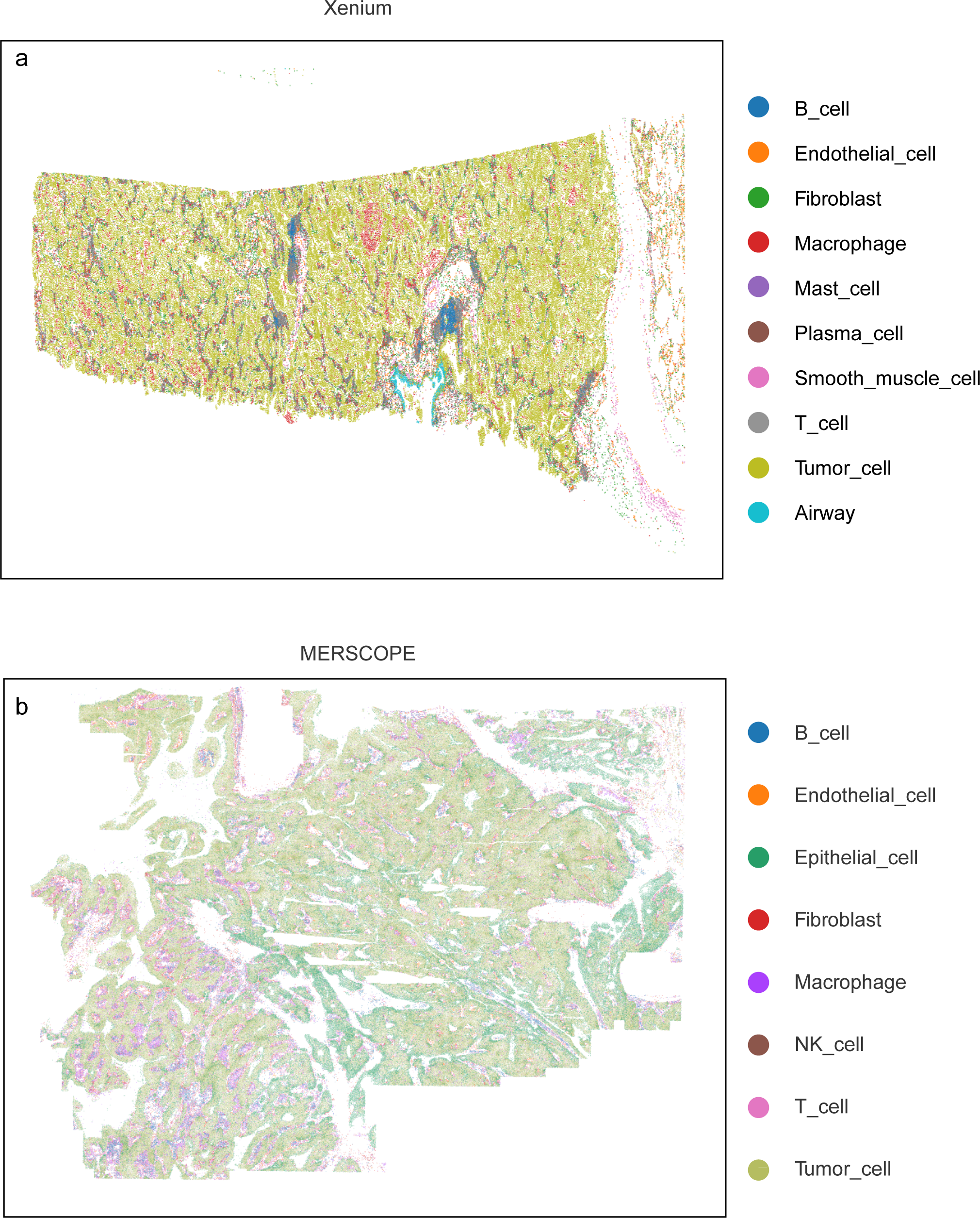
Cell type distribution in the Xenium and MERSCOPE LUAD scSRT datasets. (a) Xenium LUAD, (b) MERSCOPE LUAD. Cells are plotted based on their centroid coordinates, with colors indicating cell types.

**Fig. S2.**
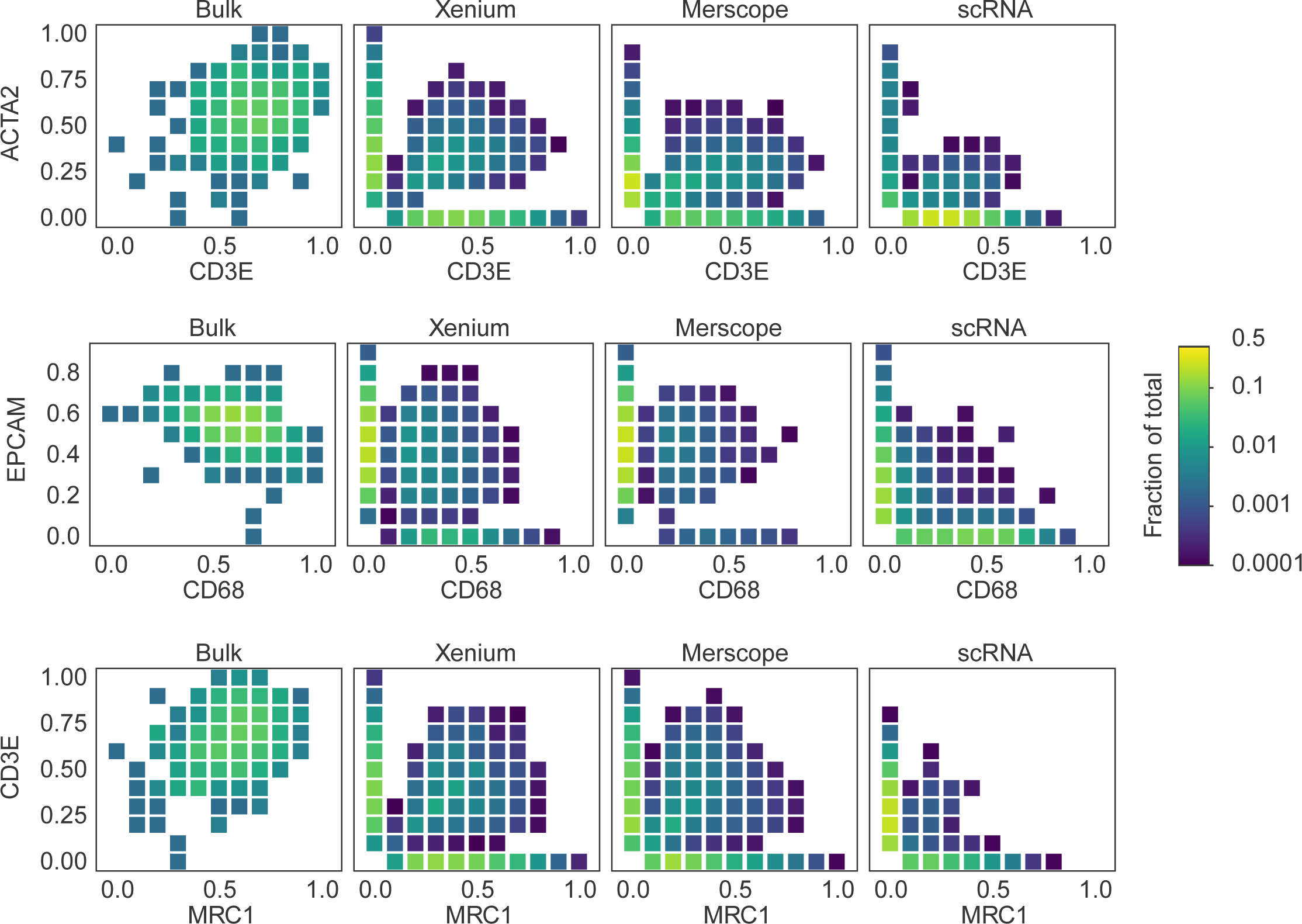
Scatter heatmaps showing mutually exclusive expression patterns for selected gene pairs. (a) ACTA2 (fibroblasts) vs CD3E (T cells); (b) EPCAM (epithelial cells) vs CD68 (macrophages); (c) CD3E (T cells) vs MRC1 (macrophages). Only cells expressing at least one gene in each pair are included. Each box represents an expression bin defined by the respective gene expression levels, with color intensity mapped logarithmically to the fraction of cells in that bin.

**Fig. S3.**
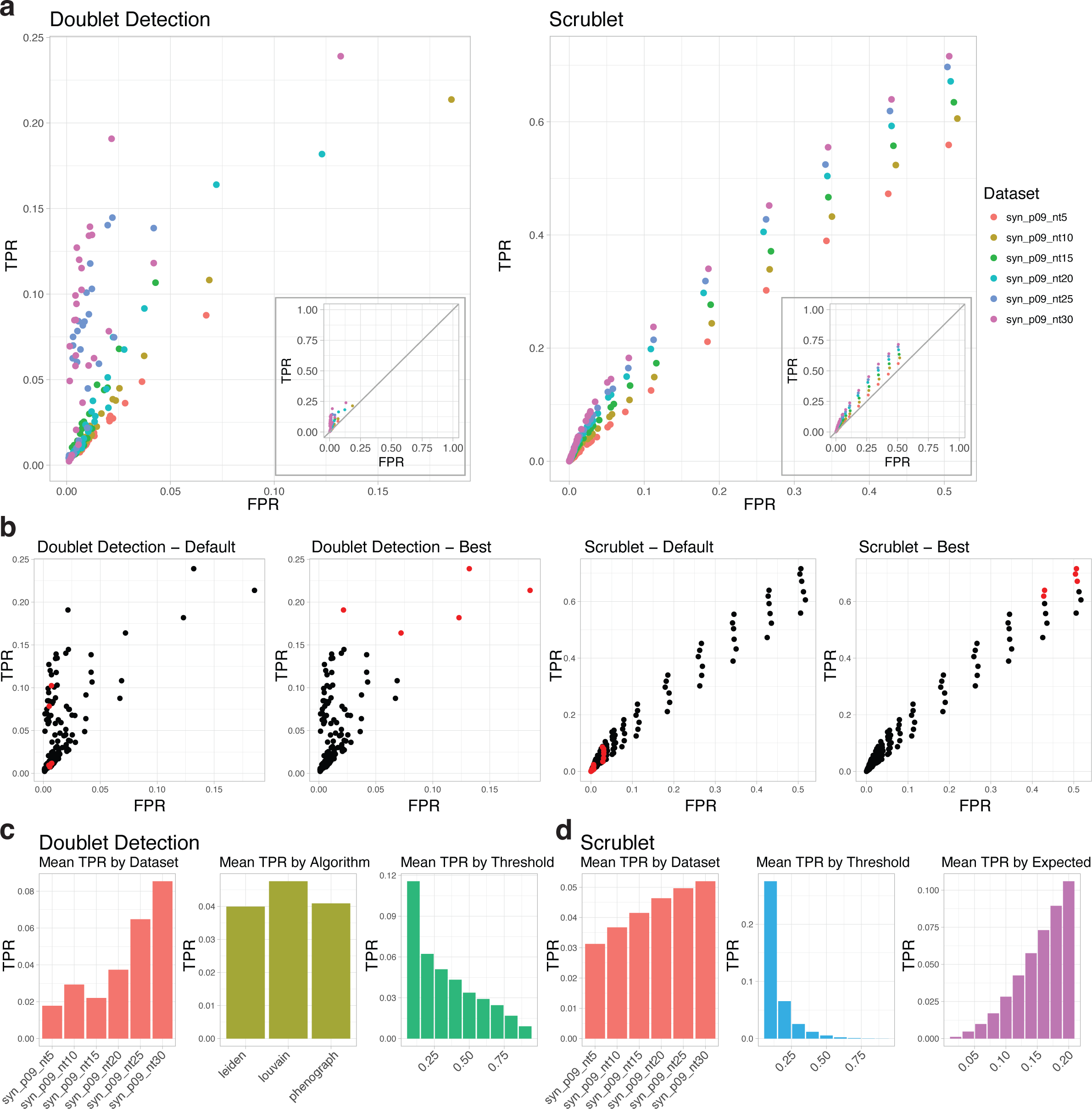
Benchmarking Traditional Doublet Detection Algorithms in Simulated Datasets. (a) False positive rate (FPR)-true positive rate (TPR) plots for DoubletDetection (left) and Scrublet (right), colored by dataset. Inset plots on 1-to-1 x- and y-axis scale, positioned so as to not cover any data points on the larger graph. (b) Results from tests using default values for DoubletDetection (louvain algorithm, 0.5 voter threshold) and the top five best performing results (highest F_1_ score) for each of the six datasets are shown in the left side plots colored in red. Results from tests using default values for Scrublet (0.06 expected doublet rate) and top five best performing results are shown in the right side plots colored in red. (c) Bar graphs showing mean TPR for DoubletDetection when using differing datasets and parameters. From left to right: dataset, algorithm selection, and voter threshold. (d) Bar graphs showing mean TPR for Scrublet. From left to right: dataset, threshold, and expected doublet rate.

**Fig. S4.**
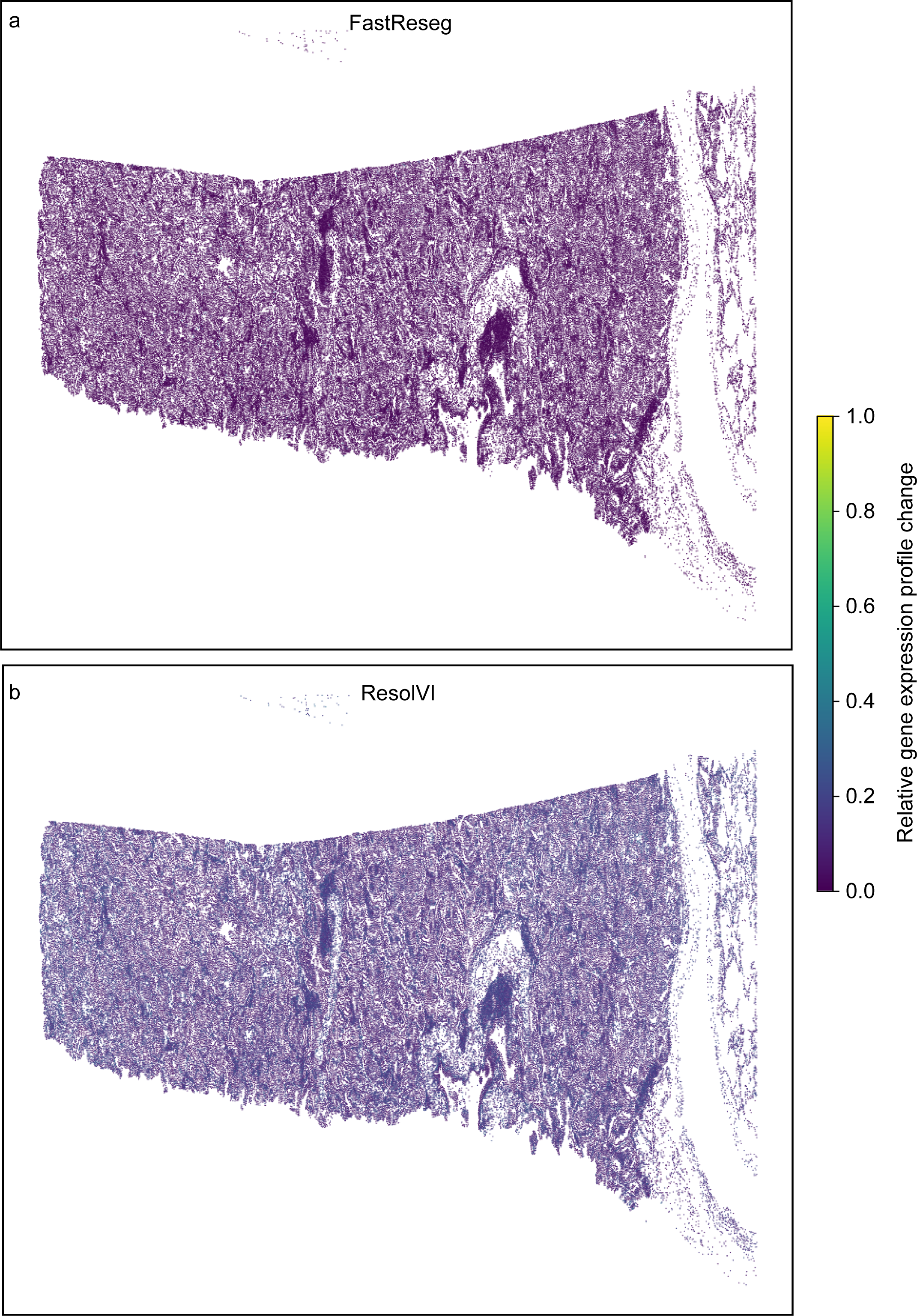
Scatter heatmap showing genes expressed exclusively in cells. Only cells expressing at least one gene in each pair are included. Each box represents an expression bin defined by the expression level of one gene in the pair, with bin colors scaled logarithmically to the fraction of cells.

**Fig. S5.**
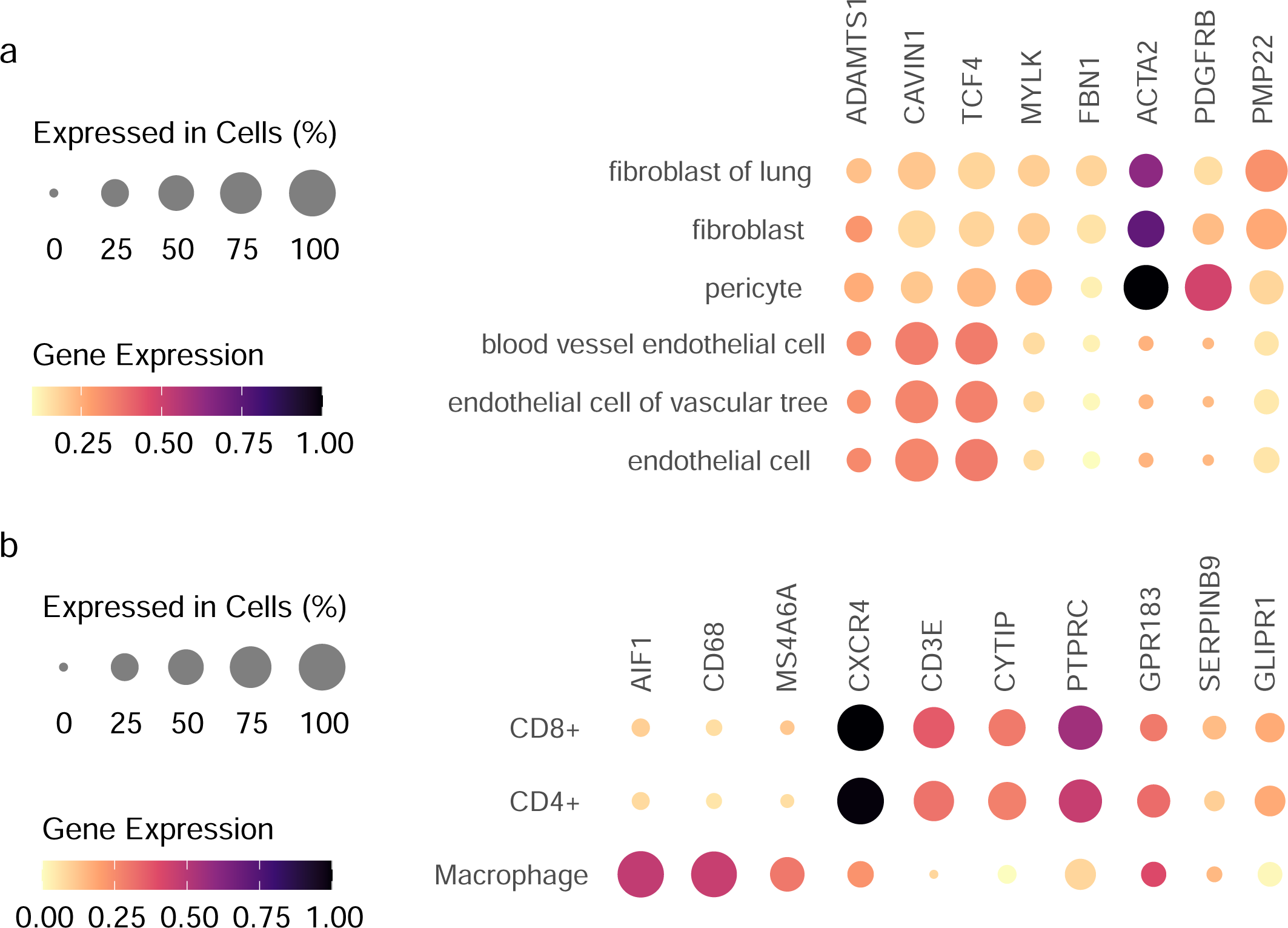
Enrichment of contaminant transcripts before MisTIC correction. Dot plots showing enrichment of contaminant transcripts among DEGs identified from uncorrected data, validated against LUAD scRNA-seq datasets. (a) Fibroblast vs. endothelial cell comparison. (b) Macrophage vs. T cell comparison. Dot size represents the percentage of cells expressing each gene; color intensity indicates average expression level.

**Fig. S6.**
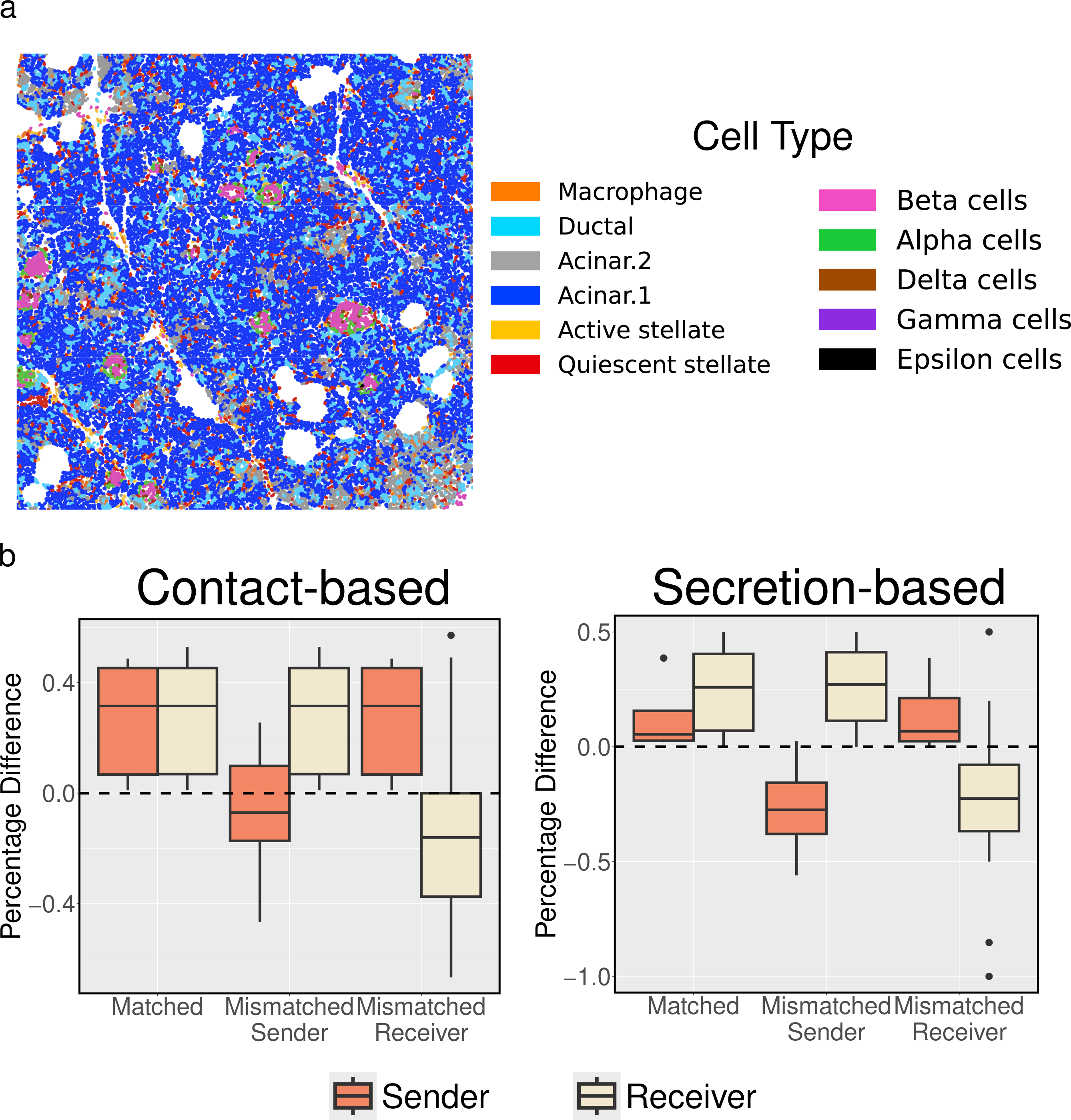
Cell-cell communication detection results on CoxMX data. (a) The CosMX pancreas SRT dataset with cell type annotation (b) Percentage changes in expression profiles by sender-receiver type with two negative control groups by contact-based CCC and secretion-based CCC.

**Table S1.** Datasets used in this study **File S1** Mathematical details of the MisTIC probabilistic model and additional analyses results

## Notes

https://github.com/yunguan-wang/MisTic-Wanglab

### Competing Interest Statement

The authors have declared no competing interest.

