## Supplementary material for "MisTIC: Missegmented Transcript Inference Correction for Improved Spatial Transcriptomics Analysis": File S1

### Supplementary File 1

#### The MisTIC probabilistic model

MisTIC is a Bayesian hierarchical model that frames the transcript misassignment correction problem as a sequence of binary classification problems. Suppose we have $t = 1, \cdots, T$ detected transcripts, $c = 1, \cdots, C$ segmented cell, $g = 1, \cdots, G$ genes, and $m = 1, \cdots, M$ cell types.

For each cell $c$, we have access to 1. $v_c$ : a vector describing its location. This contains its cell centroid and coordinates of its cell boundary polygon vertices; 2. $x_c = [x_c^1, \cdots, x_c^G]^T$ : the noisy gene counts; and 3. $\theta_c \in \{1, \cdots, M\}$: cell type assignment. For the $t$th transcript, we observe 1. $u_t \in R^2$: its spatial location; 2. $r_t \in \{1, \cdots, G\}$: its gene type; and 3. $a_t \in \{1, \cdots, C\}$: the current cell assignment based on the cell segmentation results.

Based on these quantities, MisTIC considers for each transcript $t$ whether or not it should be reassigned to its neighboring cell of a different cell type than the cell it is currently assigned to, namely, $a_t$. To simultaneously avoid implausible reassignment where the target cells are not within the vicinity of the current cell and not restrict the reassignment to only the nearest neighboring cell, we adopted a restricted kNN algorithm (1) when selecting eligible candidate neighboring cells such that only cells within a certain distance and of a different cell type will be included. Throughout the paper, we set $k = 3$ and the distance threshold to 5 microns. We have made these parameters configurable for prospect users to tailor for their own datasets. However, we find the current setting performs well across different datasets. For the following derivation, we will follow this setting, although generalization to more or fewer neighbors is straightforward.

Given a cell $c$ and its associated set of neighbor cells, to further determine whether or not to reassign a transcript $t$ to one of the cells, we further compute two quantities: 1. $b_t^k \in \{1, \cdots, C\}$: the $k$th nearest neighbor cell of transcript $t$ within the set; and 2. $z_t^k = [z_t^{k1}, z_t^{k2}, z_t^{k3}]^T$: a vector of explanatory variables for computing reassignment probability with the cell type information. Note that the $k$th nearest neighbor cell of transcript $t$ is not necessarily the $k$th nearest neighbor cell of cell $c$.

The three components of $z_t^k$ are constructed based on 1. Spatial proximity: A score measuring how close the transcript is to the boundary of the neighboring cell of a different cell type under consideration; 2. Gene expression compatibility: A score reflecting how unlikely it is for the currently assigned cell type to express the transcript's gene; and 3. Neighborhood expression support: A score capturing how highly the gene is expressed in neighboring cells of a different cell type. To be more specific, for spatial proximity, we first computed for each cell, the distances from all of its transcripts to its neighbor cells. Note that if there are multiple neighbor cells, there will be multiple distances associated with one transcript. Then, we converted the distances to percentage ranks and transformed them via a logit function. For gene expression compatibility and neighborhood expression support, we first carried out a differential gene expression analysis using DESeq2 (2) for each cell type and obtained the log2 fold change and the adjusted p-value transformed by negative of logarithm with base 10 for each gene. We then multiplied these two quantities, computed percentage ranks for all genes of a cell type, and transformed the results via a logit function. The percentage ranks and logit transformation are done to ensure the model is robust against different scales. Once all $z_t^k$'s have been computed, we associated them with the reassignment indicator $\delta_t^{kl} \in \{0, 1\}$, where $l \in \{1, 2, 3\}$ via $logit(q(\delta_t^{kl} = 1)) = \beta_0^l + \beta_1^l z_t^{kl}$, where $\beta_0^l, \beta_1^l > 0$. Note that different neighbors share the same coefficients. Along with our method of constructing $z_t^k$'s, this ensures consistency of the outputs among different neighbors. In other words, for one transcript, the nearest neighbor will always be more likely to be assigned to than the rest of the neighbor cells. And given spatial proximity and neighborhood expression support, the less expressed genes will be more likely to be reassigned. Since one transcript can only be reassigned to at most one cell, the final neighbor cell to be reassigned to will be the one with the highest reassignment probability.

To estimate $\beta_0^l$ and $\beta_1^l$, we leverage $x_c$ and $\theta_c$ to provide guidance. Specifically, fixing $k$, $\beta_0^l$ and $\beta_1^l$, we can get a set of reassignment indicators $\{\delta_t^{kl}: l \in \{1, 2, 3\}, t \in \{1, \cdots, T\}\}$. Then for cell $c$, we denote by $y_c = [y_c^1, \cdots, y_c^G]^T$, the corrected gene counts obtained via

$y_c^g = x_c^g - ||\{t: \prod_{l=1}^{3} \delta_t^{kl} = 1,\ a_t = c,\ r_t = g\}|| + ||\{t: \prod_{l=1}^{3} \delta_t^{kl} = 1,\ b_t^k = c,\ r_t = g\}||$, where $||.||$ returns the cardinality of the enclosed set. Then, we associate $y_c$ with $\theta_c$ with a logistic regression: $logit(p(\theta_c = m)) = \alpha_0^m + y_c^T \alpha^m$, where $\alpha_0^m \in R$, $\alpha^m \in R^G$.

To enable joint training, we recast the problem under the paradigm of variational inference (3). We first fix $k$ and write the log-likelihood as follows:

$$\log p(\{\theta_c\}|\{v_c\},\ \{r_t, u_t\}) = \sum_{\{\delta_t^{k1}, \delta_t^{k2}, \delta_t^{k3}\}} p(\{\delta_t^{k1}, \delta_t^{k2}, \delta_t^{k3}\}|\{v_c,\ \theta_c\},\ \{r_t, u_t\}) \log \frac{p(\{\theta_c\}, \{\delta_t^{k1}, \delta_t^{k2}, \delta_t^{k3}\}|\{v_c\}, \{r_t, u_t\})}{p(\{\delta_t^{k1}, \delta_t^{k2}, \delta_t^{k3}\}|\{v_c, \theta_c\}, \{r_t, u_t\})}$$

, where for clarity, we omitted the indices. For example, $\{\theta_c\}$ should be $\{\theta_c\}_{c=1}^{C}$.

Within the logarithm, the numerator can be further decomposed as

$$p(\{\theta_c\},\ \{\delta_t^{k1}, \delta_t^{k2}, \delta_t^{k3}\}|\{v_c\},\ \{r_t, u_t\}) = p(\{\theta_c\}|\ \{\delta_t^{k1}, \delta_t^{k2}, \delta_t^{k3}\}, \{v_c\},\ \{r_t, u_t\}) \prod_{l=1}^{3} p(\{\delta_t^{kl}\}|\{v_c\},\ \{r_t, u_t\}),$$

where we assume *a priori* independence. To compute the prior distributions, we constructed $w_t \in R^3$: a vector of explanatory variables for computing reassignment probability without the cell type information. The overall method is akin to that of $z_t^k$ with the differences being 1. the restricted kNN is computed by relaxing the cell typing constraint; and 2. the log2 fold change for each gene within a cell is computed by contrasting its neighbor cells in the expression space versus overall expressions. Throughout the paper, 50 neighbor cells were used. The same manipulation involving percentage ranks and logit transformation is applied. To generate the prior reassignment probability, the corresponding coefficients linking $w_t$ to $p(\{\delta_t^{kl}\}|\{v_c\},\ \{r_t, u_t\})$ are set by two prior beliefs. For a given transcript, if percentage ranks for two of the three components of $w_t$ are ranked top but the third is only 50%, then *a priori*, the reassignment probability is 0.01. On the other hand, if the third percentage rank is top 5%, then *a priori*, the reassignment probability is 0.99. This prior enforces the model to only reassign a transcript when three features all strongly indicate misassignment.

For the denominator, since the true posterior is unknown, we approximate it via a variational distribution $\prod_{l=1}^{3} q(\{\delta_t^{kl}\}|\{v_c, \theta_c\},\ \{r_t, u_t\})$, where $logit(q(\delta_t^{kl} = 1)) = \beta_0^l + \beta_1^l z_t^{kl}$.

Therefore, by using the Jensen's inequality, we can write the ELBO as

$$ELBO = \sum_{\{\delta_t^{k1}, \delta_t^{k2}, \delta_t^{k3}\}} \prod_{l=1}^{3} q(\{\delta_t^{kl}\}|\{v_c, \theta_c\},\ \{r_t, u_t\}) \log \frac{p(\{\theta_c\}|\ \{\delta_t^{1n}, \delta_t^{2n}, \delta_t^{3n}\}, \{v_c\}, \{r_t, u_t\}) \prod_{l=1}^{3} p(\{\delta_t^{kl}\}|\{v_c\}, \{r_t, u_t\})}{\prod_{l=1}^{3} q(\{\delta_t^{kl}\}|\{v_c, \theta_c\}, \{r_t, u_t\})}$$

This can be decomposed into

$$\sum_{\{\delta_t^{k1}, \delta_t^{k2}, \delta_t^{k3}\}} \prod_{l=1}^{3} q(\{\delta_t^{kl}\}|\{v_c, \theta_c\},\ \{r_t, u_t\}) \log p(\{\theta_c\}|\ \{\delta_t^{1n}, \delta_t^{2n}, \delta_t^{3n}\}, \{v_c\},\ \{r_t, u_t\}) \text{ and}$$

$\sum_{\{\delta_t^{k1},\delta_t^{k2},\delta_t^{k3}\}} \prod_{l=1}^{3} q(\{\delta_t^{kl}\}|\{v_c, \theta_c\}, \{r_t, u_t\}) \log \frac{\prod_{l=1}^{3} p(\{\delta_t^{kl}\}|\{v_c\}, \{r_t, u_t\})}{\prod_{l=1}^{3} q(\{\delta_t^{kl}\}|\{v_c, \theta_c\}, \{r_t, u_t\})}$. As it is computationally infeasible to sum over all possible configurations of $\{\delta_t^{k1}, \delta_t^{k2}, \delta_t^{k3}\}$, we use monte carlo integration by sampling from the variational distribution once: $\prod_{l=1}^{3} q(\{\delta_t^{kl}\}|\{v_c, \theta_c\}, \{r_t, u_t\})$. Overall, these two terms correspond to the cross-entropy and the KL-divergence. To enable a gradient-based training, we further adopted the Gumbel softmax trick (4) to sample Bernoulli random variables from $\prod_{l=1}^{3} q(\{\delta_t^{kl}\}|\{v_c, \theta_c\}, \{r_t, u_t\})$.

Due to the large number of transcripts available in a typical imaging-based SRT data (usually ranging from tens of millions to hundreds of millions), training on an entire dataset is infeasible. Therefore, we employed a tiling strategy where the SRT dataset is split into smaller overlapping patches and further removed empty patches. When loading a patch, any cell with overlap with that patch is also loaded to avoid artificially splitting a cell at boundaries. The training process then consists of sequentially loading all generated patches as minibatches, randomly selecting $k$, and performing stochastic gradient descent (5). Since the patches overlap and cover the entire dataset, the model will capture patterns across all data.

### Sensitivity analysis

The probabilistic model of MisTIC depends on the specification of several key parameters such as the mask distance cutoff beyond which transcript reassignment is no longer considered (Mask Distance), the prior probability of reassignment when a transcript is ranked top 5% given the other two factors are ranked top (Prior5), and the prior probability of reassignment when a transcript is ranked top 50% given the other two factors are ranked top (Prior50). The default values are 5µm, 99%, and 1%, respectively. We investigated the sensitivity of the MisTIC results against various configurations of each of the three parameters using the synthetic dataset. As can be seen from ***Sup. File 1 Fig. 1***, all AUROCs remain relatively invariant, demonstrating the robustness of MisTIC against different specifications.

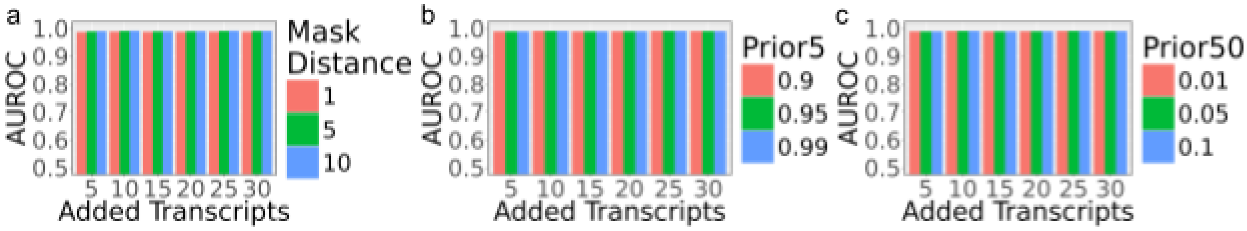

***Sup. File 1 Fig. 1*** *Sensitivity analysis on key parameters of MisTIC. (a). AUROCs by varying the mask distance; (b). AUROCs by varying the prior probability of reassignment of a transcript ranked top 5%; (c). AUROCs by varying the prior probability of reassignment of a transcript ranked top 50%.*

### Unsupervised MisTIC

Although the computation of the objective function of MisTIC relies on the cell type information, we next show that generating such information using leiden clustering (6), a widely used unsupervised clustering algorithm, can still yield comparable results using the synthetic dataset. To be more specific, we varied the resolution of the clustering parameter which indirectly controls the number of clusters with a higher resolution corresponding to a greater number of clusters. From ***Sup. File 1 Fig. 2***, we see that under most circumstances, the results are comparable to the ones with user-specified cell type information. Interestingly, when the contamination level is high (number of added transcripts ≥25), a higher resolution (a greater number of clusters) is detrimental to transcript reassignment. This suggests that overclustering would potentially create spurious reassignment.

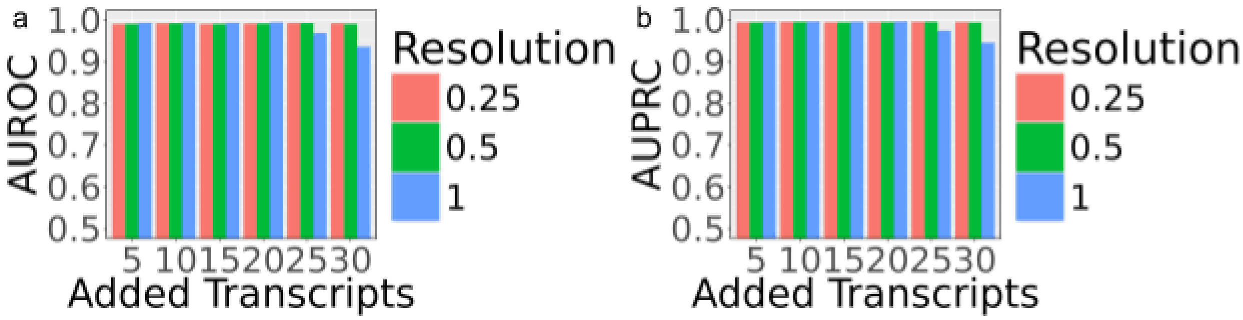


***Sup. File 1 Fig. 2*** *Analysis on MisTIC without cell type information. (a). AUROCs by varying the leiden resolution; (b). AUPRCs by varying the leiden resolution.*

### Additional benchmark results on FastReseg

In ***Fig. 2g***, we benchmarked the performance of MisTIC against the default setting of FastReseg (7). Here, we provide additional results on FastReseg on all synthetic datasets with additional configurations of its key parameters: negative decision values, gamma, and cost. Specifically, we varied the SVM class score cutoff from -2.3 to -1.7 with an increment of 0.1, changed the gamma from 0.2 to 0.4 with an increment of 0.1, and set the cost at 1, 10, and 100. As we can see from ***Sup. File 1 Fig. 3***, although different configurations have some impact on the reassignment results, they consistently underperform MisTIC by a large margin.

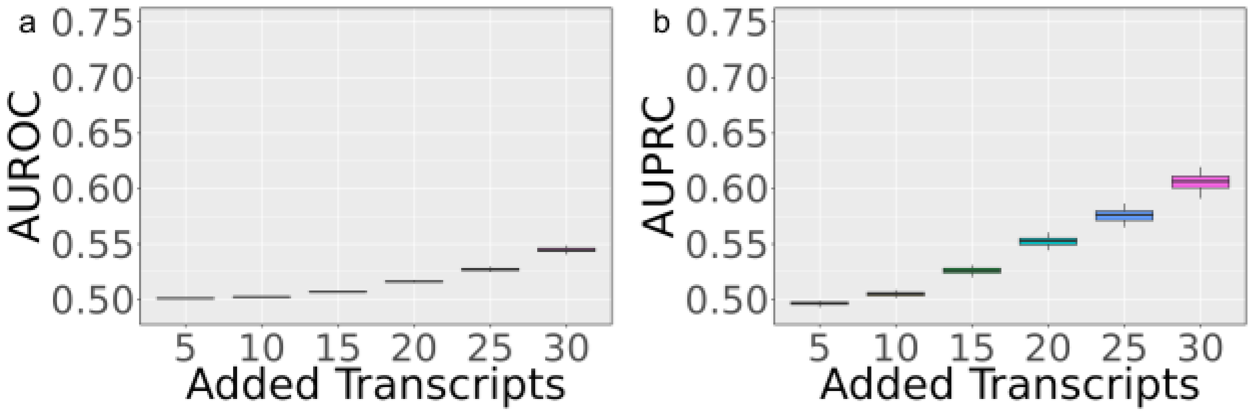


***Sup. File 1 Fig. 3*** *Additional benchmark results on FastReseg (a). AUROC; (b). AUPRC.*
