## Supplementary material for "MisTIC: Missegmented Transcript Inference Correction for Improved Spatial Transcriptomics Analysis": Table S1

| Platform | Disease | Type | Data link |
|---|---|---|---|
| Xenium | LUAD | SRT | https://www.10xgenomics.com/datasets/ffpe-human-lung-cancer-data-with-human-immuno-oncology-profiling-panel-and-custom-add-on-1-standard |
| MERSCOPE | LUAD | SRT | https://console.cloud.google.com/storage/browser/vz-ffpe-showcase/HumanLungCancerPatient1;tab=objects?pageState=(%22StorageObjectListTable%22:(%22f%22:%22%255B%255D%22))&prefix=&forceOnObjectsSortingFiltering=false |
| MERSCOPE | HCC | SRT | https://console.cloud.google.com/storage/browser/vz-ffpe-showcase/HumanLungCancerPatient1;tab=objects?pageState=(%22StorageObjectListTable%22:(%22f%22:%22%255B%255D%22))&prefix=&forceOnObjectsSortingFiltering=false |
| MERSCOPE | Prostate Cancer | SRT | https://console.cloud.google.com/storage/browser/vz-ffpe-showcase/HumanProstateCancerPatient1;tab=objects?pageState=(%22StorageObjectListTable%22:(%22f%22:%22%255B%255D%22))&prefix=&forceOnObjectsSortingFiltering=false |
| CosMX | Pancreas | SRT | https://nanostring.com/products/cosmx-spatial-molecular-imager/ffpe-dataset/cosmx-smi-human-pancreas-ffpe-dataset/ |
| Xenium Prime | Melanoma | SRT | https://www.10xgenomics.com/datasets/xenium-prime-ffpe-human-skin |
| Xenium Prime | Prostate Cancer | SRT | https://www.10xgenomics.com/datasets/xenium-prime-ffpe-human-prostate |
| TCGA | LUAD | Bulk RNA-seq | https://www.cancer.gov/ccg/research/genome-sequencing/tcga |
| scRNA-seq | LUAD | scRNA-seq | https://www.ncbi.nlm.nih.gov/geo/query/acc.cgi?acc=GSE97168 |
